# Predicting tissue-specific gene expression from whole blood transcriptome

**DOI:** 10.1101/2020.05.10.086942

**Authors:** Mahashweta Basu, Kun Wang, Eytan Ruppin, Sridhar Hannenhalli

**Affiliations:** Institute for Genome Sciences, University of Maryland, Baltimore, MD; Cancer Data Science Lab, National Cancer Institute, NIH, Bethesda, MD

**Keywords:** GTEx, Whole - Blood transcriptome, Gene expression, imputation, SNP, systems biology

## Abstract

Complex diseases are systemic, largely mediated via transcriptional dysregulation in multiple tissues. Thus, knowledge of tissue-specific transcriptome in an individual can provide important information about an individual’s health. Unfortunately, with a few exceptions such as blood, skin, and muscle, an individual’s tissue-specific transcriptome is not accessible through non-invasive means. However, due to shared genetics and regulatory programs between tissues, the transcriptome in blood may be predictive of those in other tissues, at least to some extent. Here, based on GTEx data, we address this question in a rigorous, systematic manner, for the first time. We find that an individual’s whole blood gene expression and splicing profile can predict tissue-specific expression levels in a significant manner (beyond demographic variables) for many genes. On average, across 32 tissues, the expression of about 60% of the genes is predictable from blood expression in a significant manner, with a maximum of 81% of the genes for the musculoskeletal tissue. Remarkably, the tissue-specific expression inferred from the blood transcriptome is almost as good as the actual measured tissue expression in predicting disease state for six different complex disorders, including Hypertension and Type 2 diabetes, substantially surpassing predictors built directly from the blood transcriptome. The code for our pipeline for tissue-specific gene expression prediction – TEEBoT, is provided, enabling others to study its potential translational value in other indications.

## Introduction

Most common complex (non-Mendelian) diseases involve dysfunction in multiple tissues organs. For instance, Hypertension, which is characterized by elevated arterial pressure, involves metabolic changes in heart, blood vessels, brain, kidney, etc.(Dai et al., 2018). Furthermore, the tissue and organ dysfunction is primarily driven by and reflected in transcriptional changes in various tissues and organs (Cookson, Liang, Abecasis, Moffatt, & Lathrop, 2009). Indeed, transcriptional variance mediates, in large part, causal links between genotype and complex traits (Nica et al., 2010). Thus the knowledge of tissue-specific gene expression profile can lead to a better understanding of diseases etiology, enabling patient subtyping and assessing drug efficacy (Basu et al., 2017; Grant, Pickard, Briskin, & Gutierrez-Ramos, 2002). However, except for easily accessible tissues such as blood, muscle, skin, etc. and in some cases, biopsies, organ and tissue-specific gene expression profiles cannot be readily obtained, presenting a challenge for transcription-based investigation of complex diseases. This limitation naturally gives rise to two important research questions: (1) to what extent can we predict an individual’s tissue-specific gene expression based solely on his/her whole blood gene expression, and (2) can the predicted tissue-specific expression reflect disease states better than what can be gleaned on those directly from the whole blood gene expression?

Cross-individual variance in tissue-specific gene expression can be explained, to some extent, by the genotypic variance, forming the basis of expression Quantitative Trait Loci (eQTL). A widely used tool – PrediXcan predicts tissue-specific gene expression of a gene based on the gene’s eQTL SNP (or eSNPs) (Gamazon et al., 2015). However, *a priori,* such a tool has limited scope, as it can only predict expression of a minority of genes (about 10% genes on average across tissue) that have significant tissue-specific eSNPs (Gamazon et al., 2015). Furthermore, the previous prediction models based on blood expression (e.g., (J. Wang et al., 2016) and (Halloran et al., 2015)) do not use the expression of other genes to predict the expression of a given gene, missing on the possibility of exploiting potentially shared regulatory programs between tissues. In difference, here we use an individual’s whole blood gene expression, including the whole blood splicing profile, to predict the expression of each gene in a particular tissue of the individual.

We proceed by building machine learning predictors based on GTEx data (Aguet et al., 2017) for 32 primary tissues having at least 65 samples each. We find that an individual’s whole blood gene expression and splicing profile significantly inform tissue-specific expression levels (above and beyond various demographic variable) for substantive fractions of genes, with a mean of 59% of genes across 32 tissues, up to 81% for Muscle-Skeletal tissue, based on likelihood ratio test with FDR threshold of 5%. The splicing profile contributes further beyond the gene expression profile, and for the subset of genes having eSNPs, genotype makes further significant contributions. We find that the genes with highly predictable expression are not biased toward housekeeping genes, proportionally representing tissue-specific genes. Moreover, such genes tend to have a greater number of protein interaction partners, which may suggest a contribution by shared gene networks toward expression predictability. Finally, in many cases the predicted tissue-specific expression can be used as a surrogate for actual expression in predicting disease state for several complex diseases far better than using the whole blood expression. Overall, our work establishes the utility and limits of whole blood transcriptome in estimating gene expression in other tissues, leading a basis for future translationally motivated applications. We provide our software pipeline for predicting tissue-specific gene expression, named TEEBoT (Tissue Expression Estimation using Blood Transcriptome), in a user friendly and publicly available form.

## Results

### Overview of TEEBoT – a pipeline for tissue-specific gene expression prediction

Fig. 1A illustrates the overall motivation and TEEBoT pipeline. Our goal is to assess the predictability of tissue-specific gene expression (TSGE) using all available and easily accessible information about the individual, which includes *Whole Blood* transcriptome (WBT), his/her genotype, as well as basic demographic information on age, gender and race. Normalized expression data was obtained from GTEx v6 (“The Genotype-Tissue Expression (GTEx) project.,” 2013) for 32 primary tissues (60% of all tissues) for which both the gene expression as well as WBT was available for at least 65 individuals (Supplementary Fig. S1).

**Figure 1:**
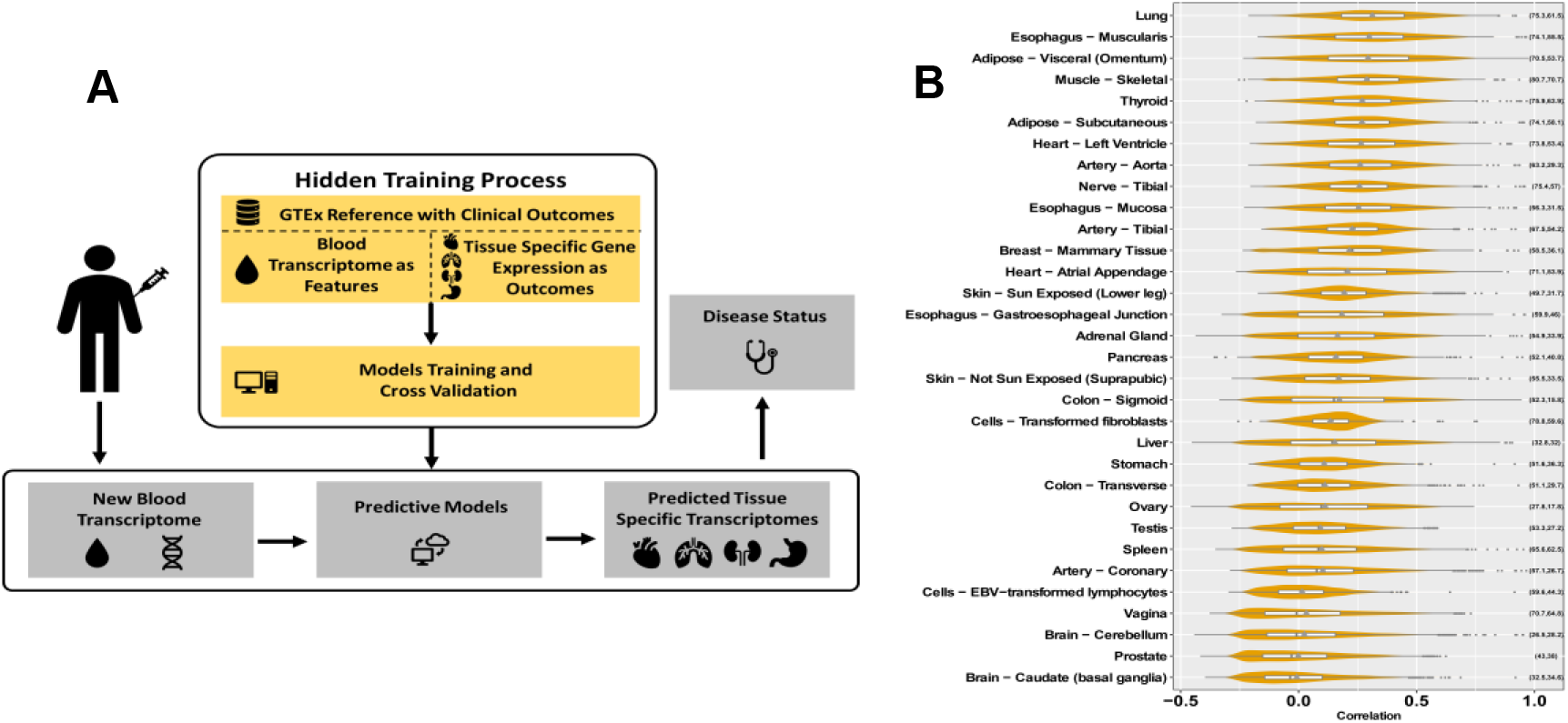
**(A)** Overall approach. We train a generalized linear model to predict tissue-specific genes expression given the Whole Blood transcriptome of an individual, which can then be used to predict disease state. **(B)** Prediction accuracy (in terms of PCC) of gene expression in target tissues from the blood expression using model M2 (WBGE+WBSp+CF), for genes with LLR (M2 ~ CF, FDR <= 0.05). The blue points mark the mean values, and the fractions of genes with significant contribution from transcriptome towards prediction over CF (LLR FDR <= 0.05) are indicated on the right side of each violin plot.?

For each tissue, and for each gene, we have fit three nested regression models to estimate TSGE: the prime model (**M2**; Methods), whose results are described in the main text, is based on Whole blood gene expression (WBGE), Whole Blood splicing (WBSp) information and three demographic ‘confounding’ factors (CF) – Age, Race, and Sex. To reduce the dimensionality of the modeling task, instead of using the expression (respectively, splicing) levels of all genes in blood, we estimated the principle components (PC) using WBGE (respectively, WBSp) across all individuals and use the sample-specific scores of top 10 PCs (top 20 PCs for WBSp) explaining 99% of variance as features. The whole-genome tissue-specific splicing profiles were obtained from (K. Wang et al., 2018). To assess the value of using splicing information we additionally built and tested our base model, ‘WBGE+CF’ model (**M1**), which uses only the WBGE PCs and CF variables. Finally, to estimate the contribution of SNPs we fit a third ‘WBGE+WBSp+SNP+CF’ model (**M3**); although this model is most inclusive it covers only a small fraction of about 10% of the genes, those having at least one eSNPs, as reported previously by the GTEx consortium (“The Genotype-Tissue Expression (GTEx) project.,” 2013). For those genes, we used the top five PCs of the genotype profile of eSNPs detected in cross-validation (CV) manner to avoid overfitting. Below we present the results obtained using our prime model M2. While results based on models M1 and M3 are mentioned briefly in context as appropriate, their details are provided as supplementary results for brevity and focus.

### The predictive power of Whole blood splicing and expression information (the M2 model)

For each of the 17,031 genes, in each of the 32 tissues we fit the regression model M2 and estimate the CV accuracies using a Pearson Correlation Coefficient (PCC) between the predicted and observed expression across individuals. First, as baseline, we assessed the contribution of WBT (i.e., WBGE and WBSp) over the demographic confounding factors via a log likelihood ratio (LLR) test and found that on average, across tissues, for 59% of genes, WBT makes a significant contribution toward TSGE prediction, with a maximum of 81% of the genes in the Muscle-Skeletal tissue; the fraction of such predicted genes and their prediction accuracies are shown in Fig. 1B. Supplementary Fig. S2 shows the corresponding plots for all genes and Supplementary Table S1 shows the number of genes with accuracy above various thresholds. For instance, on average for ~3000 (18%) of the genes, WBT makes significant contribution toward their TSGE prediction (LLR FDR <= 0.05) and they have a cross-validation PCC >= 0.3, up to 6763 genes in Muscle.

A similar assessment of the base model M1 (without WBSp) is provided in Supplementary results-1 (also Supplementary Fig. S3, and Supplementary Table S2), and a direct comparison of model M2 with the model M1 is provided in Supplementary results-2, clearly establishing the contribution of WBSp in predicting TSGE above and beyond WBGE alone; for instance, on average for 43.2% of the genes, WBSp makes a significant additional contribution (Likelihood ratio test FDR <= 0.05), up to 70.7% for Muscle-Skeletal tissue.

While SNPs are expected to contribute to TSGE prediction, as mentioned earlier, SNP-based prediction is limited to ~10% of the genes that have a significant eSNP. For the subset of such genes, we have quantified the accuracy of the model M3 which additionally includes eSNPs. As expected, for ~60% of the genes on average across tissues (this corresponds to ~6% of all genes), eSNPs make significant additional contributions (Supplementary results-3 and Supplementary Fig. S4, S5). We have also compared model M2 with a SNP-only model M4 that we have constructed (comparable to a previous tool PrediXcan (Gamazon et al., 2015)) and find that overall WBT is a better predictor of TSGE than eSNPs alone (Supplementary results-4 and Supplementary Table S3). We also found that, interestingly, genes that have eSNPs exhibit greater predictability by M2, even though M2 does not include SNPs (Supplementary Results-5; Supplementary Fig. S6).

### Characteristics of genes whose TSGE is predictable by WBT (model M2)

We investigated the distinctive properties of tissue-specific predictable genes (TSPG) by the M2 model in terms of their expression breadth, evolutionary conservation, and network connectivity. In each tissue, we identified TSPG as the genes for which WBT contributed significantly relative to CF (LLR FDR <= 0.05) and were among the top 25% most predictable based on CV PCC. First, and quite strikingly, we observe that the TSPG were quite different in each tissue (Supplementary Fig. S7), with an average Jaccard index of 0.13 across all tissue pairs.

We next identified the enriched GO biological processes in each tissue for the highly predictable genes (LLR FDR <= 0.05, CV PCC >= 0.5) using DAVID (Huang et al., 2007). The TreeMap view using REVIGO (Supek, Bošnjak, Škunca, & Šmuc, 2011) for the 17 tissues with at least 5 significantly enriched terms (FDR <= 0.05) is provided in Supplementary Fig. S8a, and a combined view of GO terms in all tissues is provided in Supplementary Fig. S8b. By and large, TSPGs are enriched for numerous fundamental cellular processes, including metabolic processes, RNA processing, translation, transcription, etc. But notably, in a few cases there is an enrichment for highly tissue-specific or tissue-relevant processes, such as ‘cardiac muscle cell action potential’ in heart and ‘cell morphogenesis involved in neuron differentiation’ in Nerve. We additionally assessed whether the broadly expressed housekeeping genes (Methods) are overrepresented among the TSPG, relative to genes with tissue-specific expression. As shown in Supplementary Fig. S9, we did not observe a substantive bias, attesting to broad utility of imputing TSGE. Looking specifically at predictability of transcription factor (TF), overall, in 19 out of 32 tissues, TF were significantly more predictable (Wilcoxon test p-value <= 0.05) than other genes; the opposite is true in only 4 cases. Supplementary Table S4 lists highly predictable (PCC >= 0.5) TFs in all tissues. Interestingly, we found that in a vast majority of tissues the TSPG are evolutionarily more conserved (Fig. 2A).

**Figure 2.**
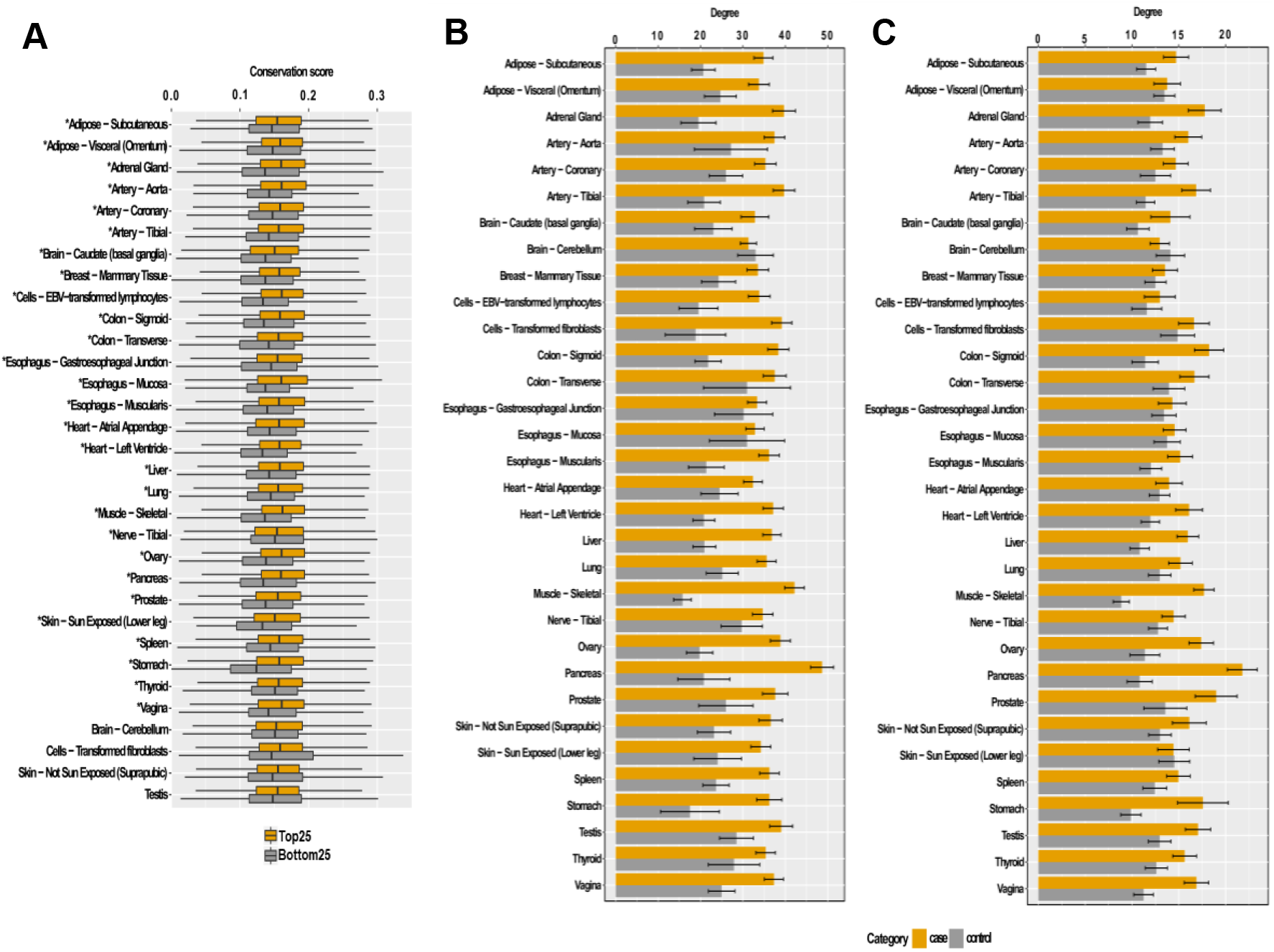
Distinguishing features of genes with predictable TSGE. **(A)** More predictable genes are more conserved. (The tissues with asterisks have significantly more conservation scores for the top25 percentile predictable genes than the bottom 25 percentile ones). The bar plots (with mean ± 95% confidence interval) shows comparison of the overall degree, **(B)** degree of housekeeping genes **(C)** of the top 25 and bottom 25 percentile predictable genes across all the tissues.

Next we assessed whether predictability of TSPG may be related to their interactions with other genes, which tend to be functionally related and have similar expression profile (Oliver, 2000; Yu et al., 2008). We therefore compared the degree distribution of TSPG in a protein interaction network (PIN) with the background (Methods). We also assessed whether broadly expressed housekeeping genes, by virtue of being expressed both in Whole Blood and the target tissue of interest, may better inform the TSGE. We therefore obtained the overall degree in the PIN as well as the degree relative only to housekeeping genes. Fig. 2B,C clearly show that TSPG have much greater connectivity, both overall, as well as relative to housekeeping genes. To further probe the potential mechanism underlying this observation, for each of the most highly predictable genes *g* in a tissue (PCC >= 0.7), we tested whether g preferentially interacts with those genes whose expression values are most predictive of g’s TSGE (Methods). We tested this hypothesis in each tissue independently using a one-sided paired Wilcoxon test across genes comparing interactions with predictive genes and the rest, using the fraction of genes in either class that interact with a given gene. We performed this test only for the tissues with at least 5 genes with non-zero interactions with the predictive features. As shown in Supplementary Table S5, in 11 of the 12 tissues in which we could test this hypothesis, it is supported (p-value <= 0.05).

For several genes, their TSGE was highly predictable (PCC >= 0.7) in multiple tissues. We investigated whether TSGE prediction model is tissue-specific by assessing whether the same or different WBT gene features were utilized in different tissues to predict the gene’s expression. For the 340 genes that are very highly predictable in multiple tissues, we estimated the overlap between the top 100 most robustly predictive features (Methods) for a given gene in two different tissues. The mean overlap between two sets of 100 features was only 8, strongly suggesting a tissue-specific model. To further probe the mechanism underlying the model’s tissue-specificity, we assessed whether the tissue-specific predictive features (genes) exhibit tissue-biased expression. Consider a gene *g* that is highly predictable in say 2 tissues *T1* and *T2,* respectively by feature sets *F1* and *F2.* We tested whether a predictive gene in F1 has higher expression in T1 compared to T2. For each of the 340 cases above, for each tissue-specific predictive gene feature, we estimated the fold difference of its expression in *T1* relative to *T2*. We found fold difference >= 1.5 in 66%, >= 2 in 56%, and >= 5 in 37% of the cases, suggesting that tissue-specific prediction utilizes distinct features, in particular those with higher expression in the specific tissue.

### Utility of WBT predicted tissue specific expression in predicting complex diseases

Finally, we assessed the extent to which the predicted TSGE can reveal tissue-specific disease-associated genes and predict disease states. For each disease annotated in GTEx and for each tissue we considered the number of samples available in the tissue that were annotated as positive for the disease and those that were annotated as negative. We retained the disease-tissue pairs having at least 25 cases (positive for the disease) and 25 controls (negative for the disease) samples in the particular tissue. This resulted in 83 disease-tissue pairs, involving 5 diseases (MHHTN-Hypertension, MHT2D-Type 2 diabetes, MHHRTATT-Acute myocardial infarction, MMHRTDIS-Ischemic heart disease, MHCOPD-Chronic respiratory disease) across 30 tissues.

We first assessed the extent to which disease-associated genes, ascertained based on observed TSGE (see below), can be identified based on the predicted TSGE. For each of the 83 disease-tissue pairs, we identified a reference set of ‘disease-associated’ genes (DG) – whose tissue-specific expression was significantly different between cases and control individuals (Wilcoxon FDR <= 0.2). We then quantified the accuracy with which the predicted TSGE could distinguish DGs from the rest of the genes. As shown in Supplementary Table S6, on average across 83 cases, predicted TSGE could distinguish DGs from the other genes with an auROC of 0.6 (with 19 cases having > 0.7 auROC). In contrast WBGE failed to predict DGs (average auROC of 0.52 with 0 cases having > 0.7 auROC). The result for Hypertension-Artery Tibial pair is shown in Fig. 3A, and disease-wise summary across all tissues shown in Fig. 3B.

**Figure 3.**
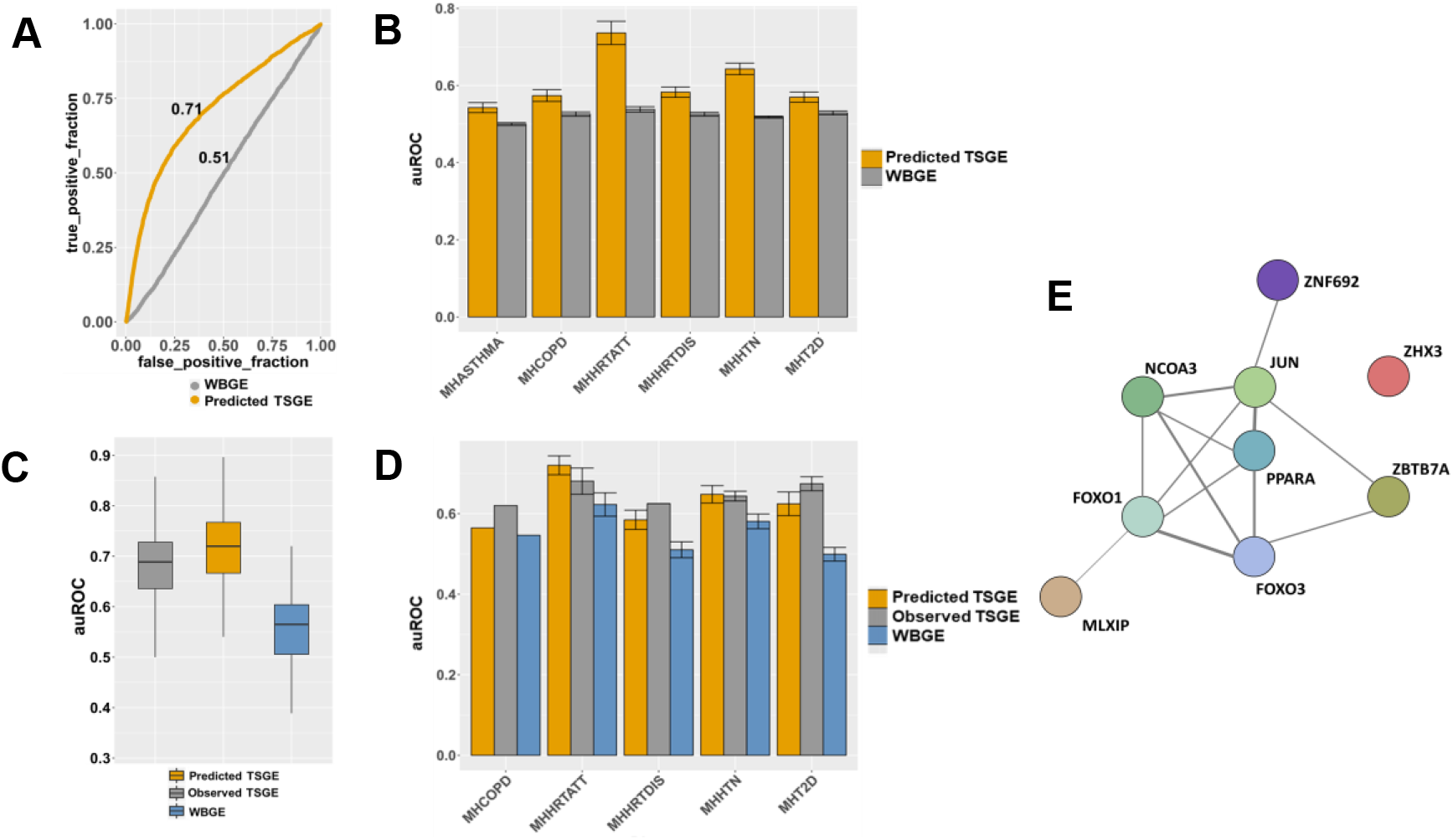
Predicting disease-genes and disease-state using predict TSGE. **(A)** ROC for prediction of genes whose observed Artery Tibial expression is associated with Hypertension based on predicted TSGE in Artery Tibial, and WBGE. **(B)** Generalization of (A) to all disease-tissue pairs: disease-wise cross-tissue summary of auROCs **(C)** Predicting Hypertension state based on observed and predicted TSGE in Artery Tibial, as well as WBGE. **(D)** Generalization of (C) to all disease-tissue pairs: disease-wise cross-tissue summary of auROCs. **(E)** Functional connection among the nine TFs that are highly predictable in Artery Tibial, and whose predicted TSGE are highly predictive of Hypertension status; see text or details.

Next, we assessed the extent to which the TSGE can predict the disease state. For this, in each disease-tissue pair, we used the genes that were highly predictable in the tissue from WBGE (LLR FDR <= 0.05 and PCC >= 0.3) as features for building the pertaining disease/control predictors. We then compared the prediction accuracy when using their (1) actual TSGE, (2) predicted TSGE, and (3) WBGE (Methods). Of the 83 disease-tissue pairs above, we focus on the 23 cases where the baseline cross-validation prediction accuracy auROC based on the actual TSGE was at least 0.6. Analysis of these 23 cases are shown in Supplementary Fig. S10 and Supplementary Table S7. Results for one specific case of Hypertension-Artery Tibial is shown in Fig. 3C, and disease-wise summary across tissues are shown in Fig. 3D. These results shown that (i) the accuracies using predicted TSGE is comparable to those using observed TSGE (average fractional difference = 0.3%), (ii) predicted TSGE performs substantially better than WBGE (average fractional difference = 12%), and (iii) WBGE performance is modest (average auROC = 0.57). Overall, these results suggest that predicted TSGE can provide insights into tissue-specific disease-linked genes, and can predict disease-state, comparable to observed TSGE, and is superior than WBGE.

We illustrate the above results for specific case of Hypertension-Artery Tibial pair (Fig. 3A,C). There are 108 genes (i) whose gene expression in Artery Tibial are highly predictable using WBT (PCC >= 0.5), and (ii) whose predicted TSGE was differential between Hypertensive individuals relative to control group (p-value <= 0.05).These genes are enriched for two major functional categories – Various acid metabolism including Carboxylic acid, and various ion and Carboxylic acid transports, both of which have functional links with Hypertension (Das, 2010; Ives, 1989). The 108 genes include 9 TFs shown in Fig. 3E, 8 of which are functionally related based on diverse array of evidence according to STRING database (Franceschini et al., 2013). PPAR-alpha and NCOA3 (also known as SRC3) form the two hubs. PPAR-alpha, by virtue of its involvement in control of vascular tone, has been suggested as an important target for Hypertension (Usuda, 2014). SRC3 is also known to regulates smooth muscle cell transcription thus regulating hypertension (Li et al., 2007). Other TFs also have links to Hypertension. FOXO1 is involved in vascular homeostasis (Dharaneeswaran et al., 2014), and FOXO3 variants have been linked to blood pressure (Morris et al., 2016). ZNF692 (also known as AREBP) is linked by GWAS to systolic blood pressure in gene cards database. ZHX3 (Yamada et al., 2018) and MLXIP (Alobeidy et al., 2013) have been linked to coronary artery disease and Hypertension. JUN (also known as AP-1) was associated with arterial stiffness in elderly Hypertensive patients (Liu et al., 2016). Overall, this example illustrates the potential clinical value of WBT-based TSGE prediction.

## Discussion

Charting tissue-specific gene expression profiles in individuals is important for understanding complex diseases, a realization that has been one of the prime motivations of the GTEx consortium (Ongen et al., 2017). Here we utilized the availability of genotypes and tissue-wise gene expression profiles in dozens of tissues across hundreds of individuals in the GTEx database to build models that predict tissue-specific gene expression (TSGE) profiles from the blood, which is by far the most readily available tissue. This particular scenario has not been comprehensively evaluated previously. Furthermore, we show, for the first time that the global splicing profile in the blood significantly contributes to the predictability of TSGE in other tissues of the same individual.

Our results show that the more predictable genes have a greater connectivity to other genes in a protein-protein interaction network. However, interestingly, such highly predictable genes are not particularly biased towards broadly expressed housekeeping genes and proportionally represent genes with tissue-restricted expression, attesting to a broad utility of the approach.

Most importantly, we find that the predicted TSGE using model M2, which learns from WBT alone, performs far better than the source WBT in predicting disease state. That is, the global expression and splicing profile in blood captures clinically relevant information indirectly via predicted TSGE in other tissues better than when it is used directly as a surrogate for TSGE.

Taken together, our results provides a comprehensive and positive response to the two research questions we have set to study: it charts the extent to which human tissue expression can be predicted from blood transcriptomics in 25 human tissues, and based on the latter, it lays a basis for the future utilization of blood expression data for building predictive models of complex disorders.

## Methods

### Linear models for gene expression predictability

We used three different linear regression models to predict a gene’s expression in a tissue (other than Whole Blood):

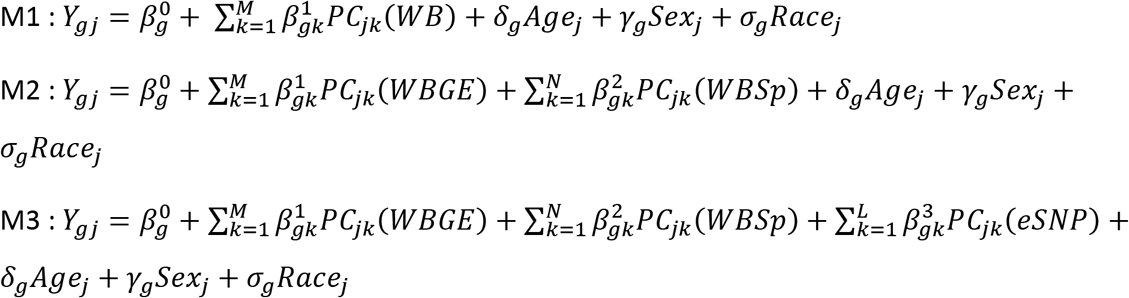

where *Y_gj_* is the expression of *g^th^* gene in the target tissue in *j^th^* sample, *PC_jk_(WBGE), PCj_k_(WBSp), and PC_jk_(eSNP)* denote the value of *k^th^* principal component (PC) of Whole Blood gene expression, Whole Blood splicing, and eSNPs for *j^th^* sample respectively. *Agej, Sexj, and Racej* denote the age, sex, and race of *j^th^* sample. Note that, instead of using all genes’ expression and splicing in WB, we use a reduced PC representation to prevent overfitting, while still capturing the variability. Specifically, we use top 10 PCs based on the WBGE, 20 PCs for WBSp across the GTEx individuals as representative WBC transcriptomic features, capturing 99% variance. To avoid overfitting, the eSNPs used in model M3 are determined in each of the cross-validation step and then the top 5 PCs of these eSNPs are used for prediction. eSNPs which are present within 1MB region of the corresponding gene were used in the model. LASSO package from R is used to build the regression model, and results are computed for 5-fold cross validation with 25 independent iterations.

In order to assess the contribution of SNPs without the WBT in prediction of gene expression we build model M4

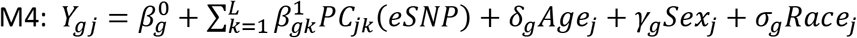

Only the eSNPs within 1MB region of the gene “g”, identified in the training set alone are considered. Apart from these models, we also implemented a baseline model M0 based only on the confounders – Age, Sex, and Race.

### Log-likelihood ratio test to identify genes whose expression is informed by various model features

For each of the three models (M1, M2 and M3) we assess for each gene whether its expression predictability has significant contribution from WBGE, (WBGE+WBSp), and (WBGE+WBSp+eSNPs) respectively, above and beyond Age, Race, and Sex. For each gene we compare the model (either M1, M2, or M3) with the Null model M0 using Log-likelihood ratio test using R package “lmtest”. The p-value indicates the significance of the contribution by the additional features. We apply FDR <= 0.05 to select the genes, henceforth called the “significant gene” with respect to a particular model. With regard to finding the “significant genes” that has significant contribution from eSNPs above and beyond WBT, we compare model M4 with M3 using the Log-likelihood ratio test as above.

### Characterizing predictable genes

#### Housekeeping genes

3791 housekeeping genes (HK) were obtained from article (Eisenberg & Levanon, 2013), of which 3342 genes were common to the GTEx gene sets and were considered. *Tissue Specificity:* For estimating tissues specificity of each gene we use GTEx data version 6. For each of the gene in a target tissue, we calculate its tissue specificity as log_2_ of ratio of mean gene expression in target tissue to the mean gene expression in rest of the tissues and consider the genes with their tissue specificity among the top 25 percentile.

#### Connectivity

The protein interaction network is obtained from (Schaefer et al., 2012). From this network we extract the degrees of connectivity of the top and bottom 25 percentile predictable genes, which are considered as foreground and control for comparison of their degrees. Later we perform a similar comparison by considering only the connectivity with the HK genes.

### Identifying most predictive features of a gene

For a given gene in a particular tissue, we find the list of genes whose expression in whole blood contribute significantly toward its expression prediction. We consider the top 5PCs (most frequently appearing across independent tuns) of blood gene expression which contribute to the prediction, and for each PC we identify top 20 genes which are most correlated with the corresponding PC. Overall this yields 100 genes (across 5 PCs), denoted as *S(g),* contributing significantly toward g’s TSGE prediction.

### Analysis of disease prediction

The disease genes are determined for each disease-tissue pair for each 5-fold cross-validation iteration, based on Wilcoxon test for each gene using its expression value across the disease and non-disease individuals among the training samples. Genes with FDR <= 0.2 are considered disease genes (DGs). For each DG we also find its disease predictability, in the test samples, using raw TSGE, blood expression and predicted TSGE using lasso in glmnet R-package. Additionally, we estimate disease predictability specific to tissue taking into account all the genes whose expression are significantly predictable (FDR LLR <=0.05 and predictability score >=0.3). To do so we build a Lasso model and estimate auROC in a cross-validation fashion.

## Supporting information

Supplemental Results

## References

Aguet, F., Ardlie, K. G., Cummings, B. B., Gelfand, E. T., Getz, G., Hadley, K., … Montgomery, S. B. (2017). Genetic effects on gene expression across human tissues. Nature, 550(7675), 204–213. https://doi.org/10.1038/nature24277

Alobeidy, B. F., Li, C., Alzobair, A. A., Liu, T., Zhao, J., Fang, Y., & Zheng, F. (2013). The Association Study between Twenty One Polymorphisms in Seven Candidate Genes and Coronary Heart Diseases in Chinese Han Population. PLoS ONE. https://doi.org/10.1371/journal.pone.0066976

Basu, M., Sharmin, M., Das, A., Nair, N. U., Wang, K., Lee, J. S., … Hannenhalli, S. (2017). Prediction and subtyping of hypertension from pan-tissue transcriptomic and genetic analyses. Genetics. https://doi.org/10.1534/genetics.117.300280

Cookson, W., Liang, L., Abecasis, G., Moffatt, M., & Lathrop, M. (2009). Mapping complex disease traits with global gene expression. Nature Reviews Genetics. https://doi.org/10.1038/nrg2537

Dai, X., Hua, L., Chen, Y., Wang, J., Li, J., Wu, F., … Liang, C. (2018). Mechanisms in hypertension and target organ damage: Is the role of the thymus key? (Review). International Journal of Molecular Medicine. https://doi.org/10.3892/ijmm.2018.3605

Das, U. N. (2010). Essential fatty acids and their metabolites in the context of hypertension. Hypertension Research. https://doi.org/10.1038/hr.2010.105

Dharaneeswaran, H., Abid, M. R., Yuan, L., Dupuis, D., Beeler, D., Spokes, K. C., … Aird, W. C. (2014). FOXO1-mediated activation of akt plays a critical role in vascular homeostasis. Circulation Research. https://doi.org/10.1161/CIRCRESAHA.115.303227

Eisenberg, E., & Levanon, E. Y. (2013). Human housekeeping genes, revisited. Trends in Genetics. https://doi.org/10.1016/j.tig.2013.05.010

Franceschini, A., Szklarczyk, D., Frankild, S., Kuhn, M., Simonovic, M., Roth, A., … Jensen, L. J. (2013). STRING v9.1: Protein-protein interaction networks, with increased coverage and integration. Nucleic Acids Research, 41(November 2012), 808–815. https://doi.org/10.1093/nar/gks1094

Gamazon, E. R., Wheeler, H. E., Shah, K. P., Mozaffari, S. V, Aquino-Michaels, K., Carroll, R. J.,… Im, H. K. (2015). A gene-based association method for mapping traits using reference transcriptome data. Nature Genetics, 47(9), 1091–1098. https://doi.org/10.1038/ng.3367

Grant, E. P., Pickard, M. D., Briskin, M. J., & Gutierrez-Ramos, J. C. (2002). Gene expression profiles: Creating new perspectives in arthritis research. Arthritis and Rheumatism. https://doi.org/10.1002/art.10014

Halloran, J. W., Zhu, D., Qian, D. C., Byun, J., Gorlova, O. Y., Amos, C. I., & Gorlov, I. P. (2015). Prediction of the gene expression in normal lung tissue by the gene expression in blood. BMC Medical Genomics. https://doi.org/10.1186/s12920-015-0152-7

Huang, D. W., Sherman, B. T., Tan, Q., Kir, J., Liu, D., Bryant, D., … Lempicki, R. A. (2007). DAVID Bioinformatics Resources: Expanded annotation database and novel algorithms to better extract biology from large gene lists. Nucleic Acids Research. https://doi.org/10.1093/nar/gkm415

Ives, H. E. (1989). Ion transport defects and hypertension where is the link? Hypertension. https://doi.org/10.1161/01.HYP.14.6.590

Li, H. J., Haque, Z., Lu, Q., Li, L., Karas, R., & Mendelsohn, M. (2007). Steroid receptor coactivator 3 is a coactivator for myocardin, the regulator of smooth muscle transcription and differentiation. Proceedings of the National Academy of Sciences of the United States of America. https://doi.org/10.1073/pnas.0611639104

Liu, Q., Han, L., Du, Q., Zhang, M., Zhou, S., & Shen, X. (2016). The association between oxidative stress, activator protein-1, inflammatory, total antioxidant status and artery stiffness and the efficacy of olmesartan in elderly patients with mild-to-moderate essential hypertension. Clinical and Experimental Hypertension. https://doi.org/10.3109/10641963.2015.1131285

Morris, B. J., Chen, R., Donlon, T. A., Evans, D. S., Tranah, G. J., Parimi, N., … Willcox, B. J. (2016). Association Analysis of FOXO3 Longevity Variants with Blood Pressure and Essential Hypertension. American Journal of Hypertension. https://doi.org/10.1093/ajh/hpv171

Nica, A. C., Montgomery, S. B., Dimas, A. S., Stranger, B. E., Beazley, C., Barroso, I., & Dermitzakis, E. T. (2010). Candidate causal regulatory effects by integration of expression QTLs with complex trait genetic associations. PLoS Genetics. https://doi.org/10.1371/journal.pgen.1000895

Oliver, S. (2000). Guilt-by-association goes global. Nature, 403(6770), 601–603. https://doi.org/10.1038/35001165

Ongen, H., Brown, A. A., Delaneau, O., Panousis, N. I., Nica, A. C., & Dermitzakis, E. T. (2017). Estimating the causal tissues for complex traits and diseases. Nature Genetics. https://doi.org/10.1038/ng.3981

Schaefer, M. H., Fontaine, J. F., Vinayagam, A., Porras, P., Wanker, E. E., & Andrade-Navarro, M. A. (2012). Hippie: Integrating protein interaction networks with experiment based quality scores. PLoS ONE, 7(2). https://doi.org/10.1371/journal.pone.0031826

Supek, F., Bošnjak, M., Škunca, N., & Šmuc, T. (2011). Revigo summarizes and visualizes long lists of gene ontology terms. PLoS ONE. https://doi.org/10.1371/journal.pone.0021800

The Genotype-Tissue Expression (GTEx) project. (2013). Nature Genetics, 45(6), 580–585. https://doi.org/10.1038/ng.2653

Usuda, D. (2014). Peroxisome proliferator-activated receptors for hypertension. World Journal of Cardiology. https://doi.org/10.4330/wjc.v6.i8.744

Wang, J., Gamazon, E. R., Pierce, B. L., Stranger, B. E., Im, H. K., Gibbons, R. D., … Chen, L. S. (2016). Imputing Gene Expression in Uncollected Tissues Within and beyond GTEx. American Journal of Human Genetics. https://doi.org/10.1016/j.ajhg.2016.02.020

Wang, K., Wu, D., Zhang, H., Das, A., Basu, M., Malin, J., … Hannenhalli, S. (2018). Comprehensive map of age-associated splicing changes across human tissues and their contributions to age-associated diseases. Scientific Reports. https://doi.org/10.1038/s41598-018-29086-2

Yamada, Y., Yasukochi, Y., Kato, K., Oguri, M., Horibe, H., Fujimaki, T., … Sakuma, J. U. N. (2018). Identification of 26 novel loci that confer susceptibility to early-onset coronary artery disease in a Japanese population. Biomedical Reports. https://doi.org/10.3892/br.2018.1152

Yu, H., Braun, P., Yildirim, M. A., Lemmens, I., Venkatesan, K., Sahalie, J., … Vidal, M. (2008). High-quality binary protein interaction map of the yeast interactome network. Science. https://doi.org/10.1126/science.1158684

