## Supplementary figures and images for "Predicting tissue-specific gene expression from whole blood transcriptome"

### Supplemental Figure S8b

ALLGO Gene Ontology treemap

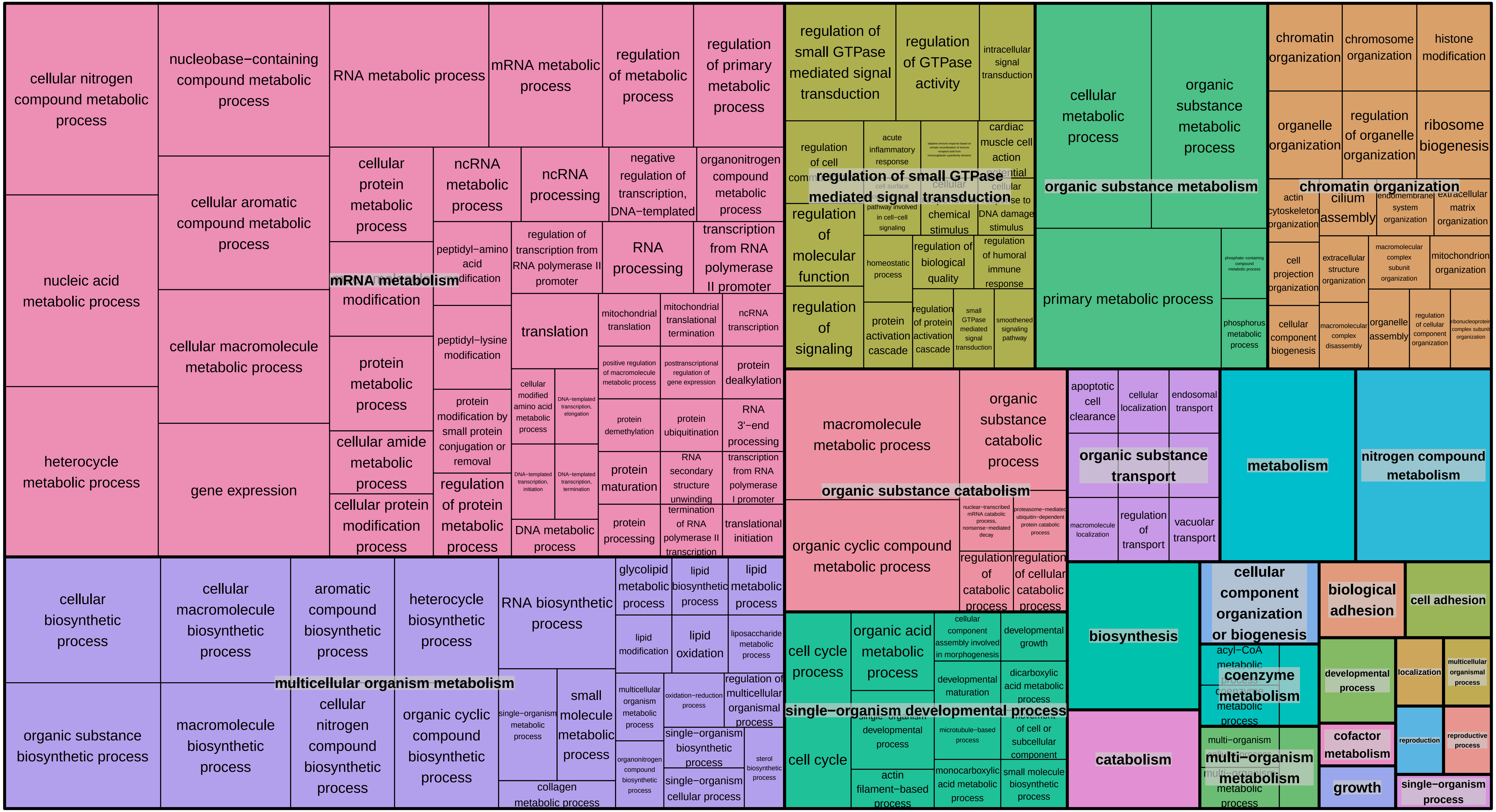
