## Supplemental Figure S8a for "Predicting tissue-specific gene expression from whole blood transcriptome"

### Adipose−Subcutaneous Gene Ontology treemap

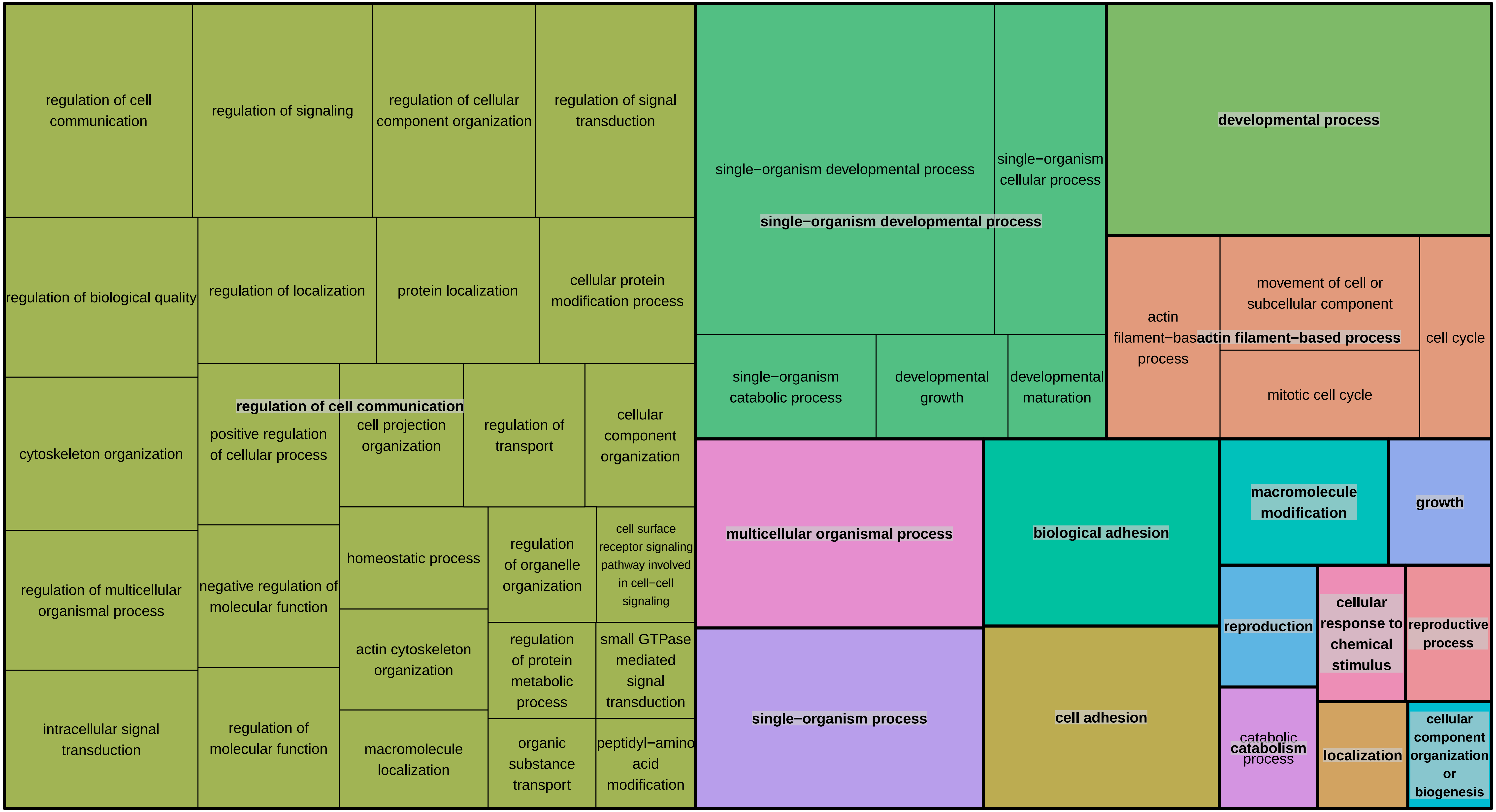

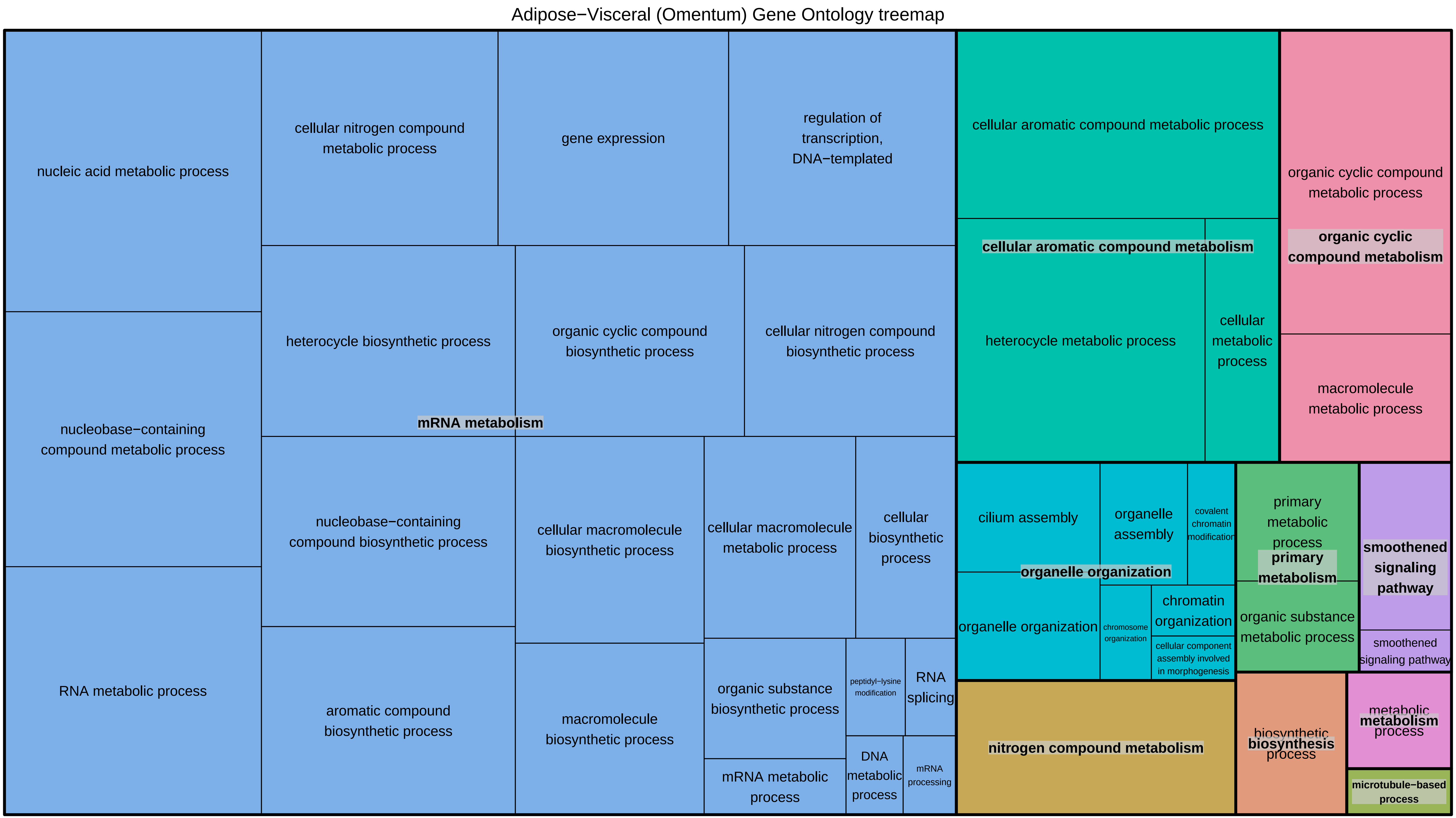
Adipose−Visceral (Omentum) Gene Ontology treemap
nucleic acid metabolic process
cellular nitrogen compound metabolic process
gene expression
regulation of transcription, DNA−templated
cellular aromatic compound metabolic process
organic cyclic compound metabolic process
organic cyclic compound metabolism
cellular aromatic compound metabolism
heterocycle biosynthetic process
organic cyclic compound biosynthetic process
cellular nitrogen compound biosynthetic process
heterocycle metabolic process
cellular metabolic process
macromolecule metabolic process
nucleobase−containing compound metabolic process
mRNA metabolism
nucleobase−containing compound biosynthetic process
cellular macromolecule biosynthetic process
cellular macromolecule metabolic process
cellular biosynthetic process
cilium assembly
organelle assembly
covalent chromatin modification
primary metabolic process
primary metabolism
smoothened signaling pathway
organelle organization
organelle organization
chromosome organization
chromatin organization
cellular component assembly involved in morphogenesis
organic substance metabolic process
smoothened signaling pathway
RNA metabolic process
aromatic compound biosynthetic process
macromolecule biosynthetic process
organic substance biosynthetic process
peptidyl−lysine modification
RNA splicing
nitrogen compound metabolism
biosynthetic process
biosynthesis
metabolic process
metabolism
mRNA metabolic process
DNA metabolic process
mRNA processing
microtubule−based process

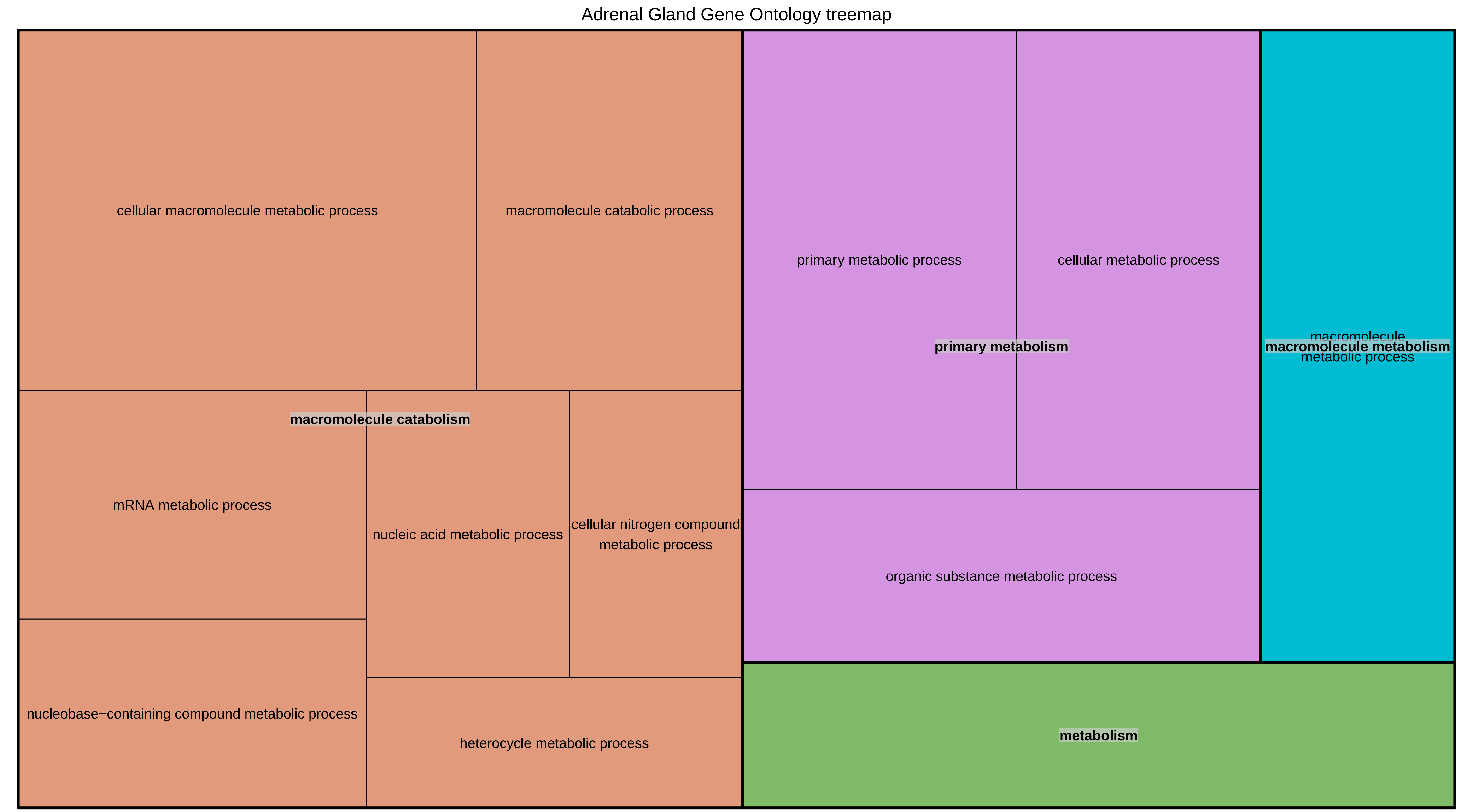
Adrenal Gland Gene Ontology treemap
cellular macromolecule metabolic process
macromolecule catabolic process
macromolecule catabolism
mRNA metabolic process
nucleic acid metabolic process
cellular nitrogen compound metabolic process
nucleobase−containing compound metabolic process
heterocycle metabolic process
primary metabolic process
cellular metabolic process
primary metabolism
organic substance metabolic process
macromolecule metabolic process
macromolecule metabolism
metabolism

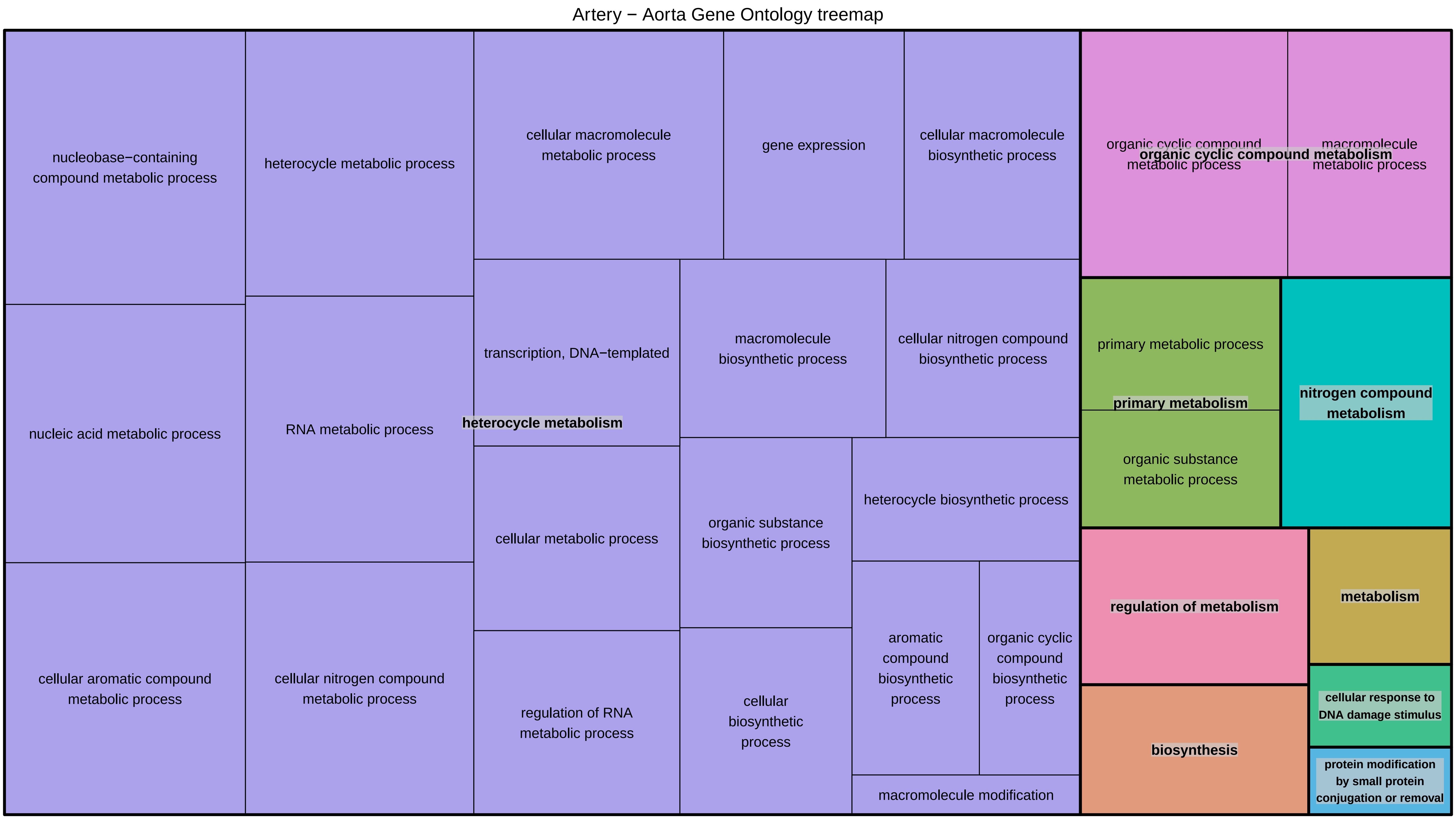
Artery − Aorta Gene Ontology treemap
nucleobase−containing compound metabolic process
heterocycle metabolic process
cellular macromolecule metabolic process
gene expression
cellular macromolecule biosynthetic process
organic cyclic compound metabolic process
macromolecule metabolic process
organic cyclic compound metabolism
transcription, DNA−templated
macromolecule biosynthetic process
cellular nitrogen compound biosynthetic process
primary metabolic process
primary metabolism
nitrogen compound metabolism
nucleic acid metabolic process
RNA metabolic process
heterocycle metabolism
organic substance metabolic process
cellular metabolic process
organic substance biosynthetic process
heterocycle biosynthetic process
regulation of metabolism
metabolism
cellular aromatic compound metabolic process
cellular nitrogen compound metabolic process
regulation of RNA metabolic process
cellular biosynthetic process
aromatic compound biosynthetic process
organic cyclic compound biosynthetic process
cellular response to DNA damage stimulus
biosynthesis
protein modification by small protein conjugation or removal
macromolecule modification

Artery − Coronary Gene Ontology treemap

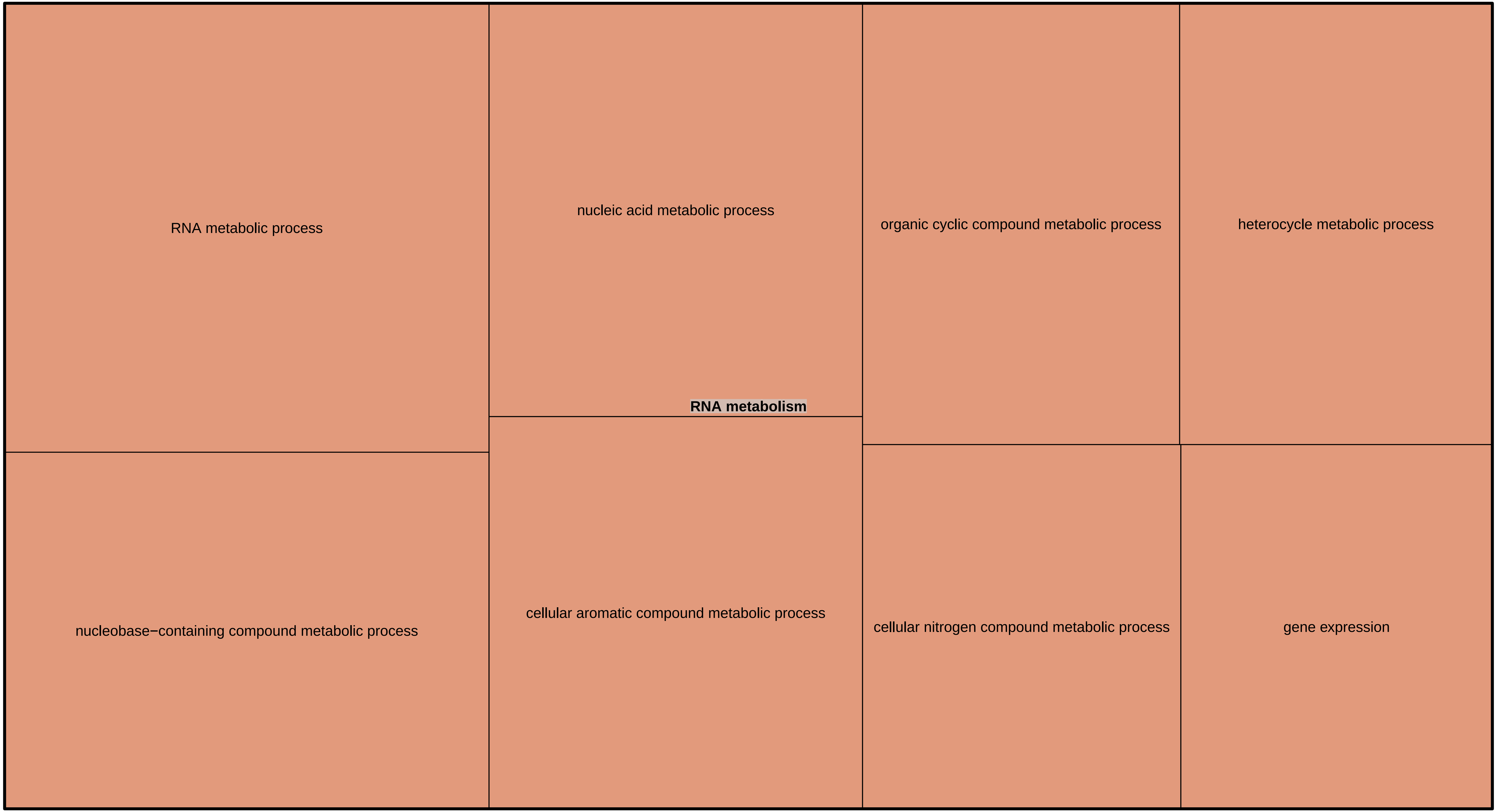

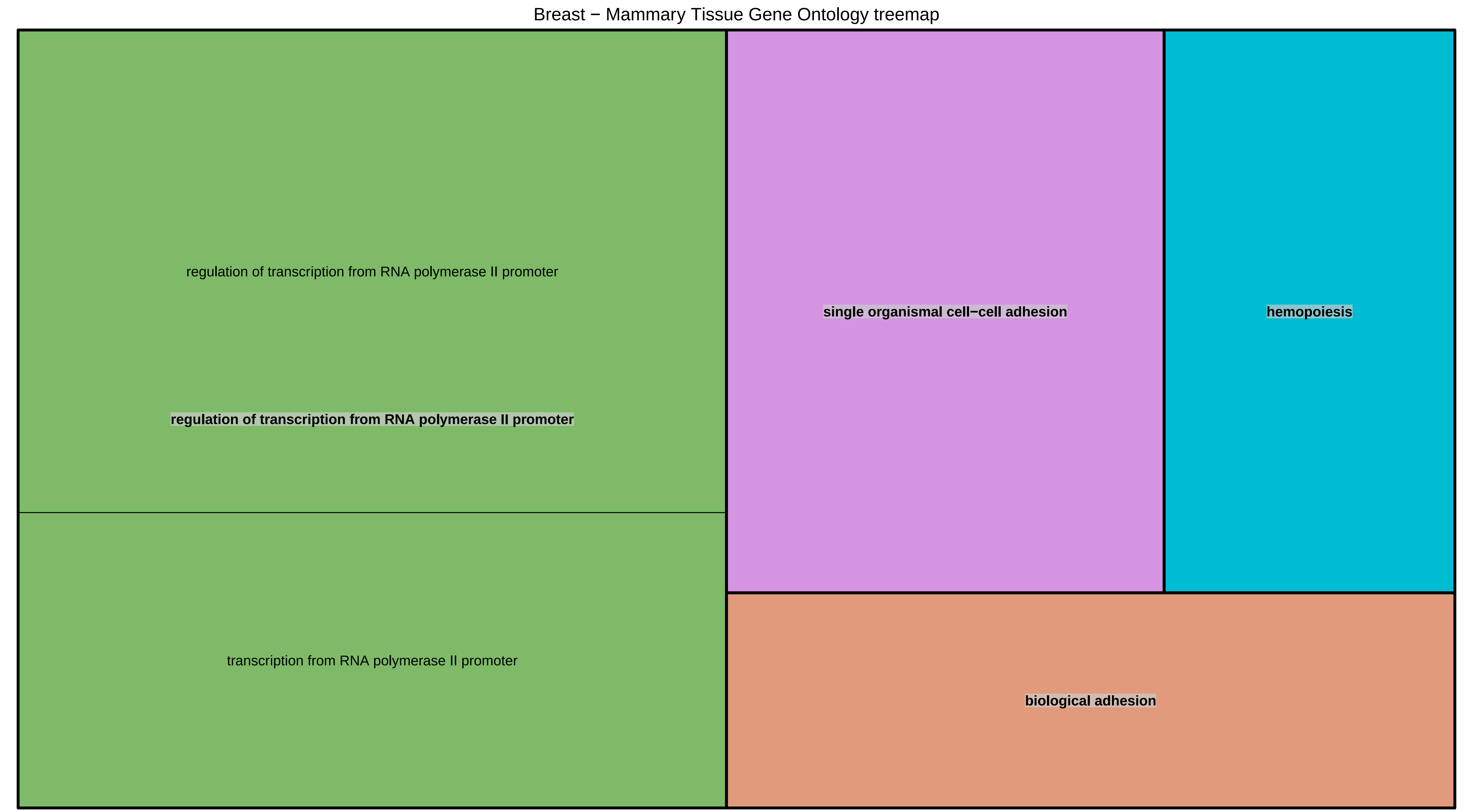
Breast − Mammary Tissue Gene Ontology treemap
regulation of transcription from RNA polymerase II promoter
regulation of transcription from RNA polymerase II promoter
transcription from RNA polymerase II promoter
single organismal cell−cell adhesion
hemopoiesis
biological adhesion

Esophagus − Gastroesophageal Junction Gene Ontology treemap

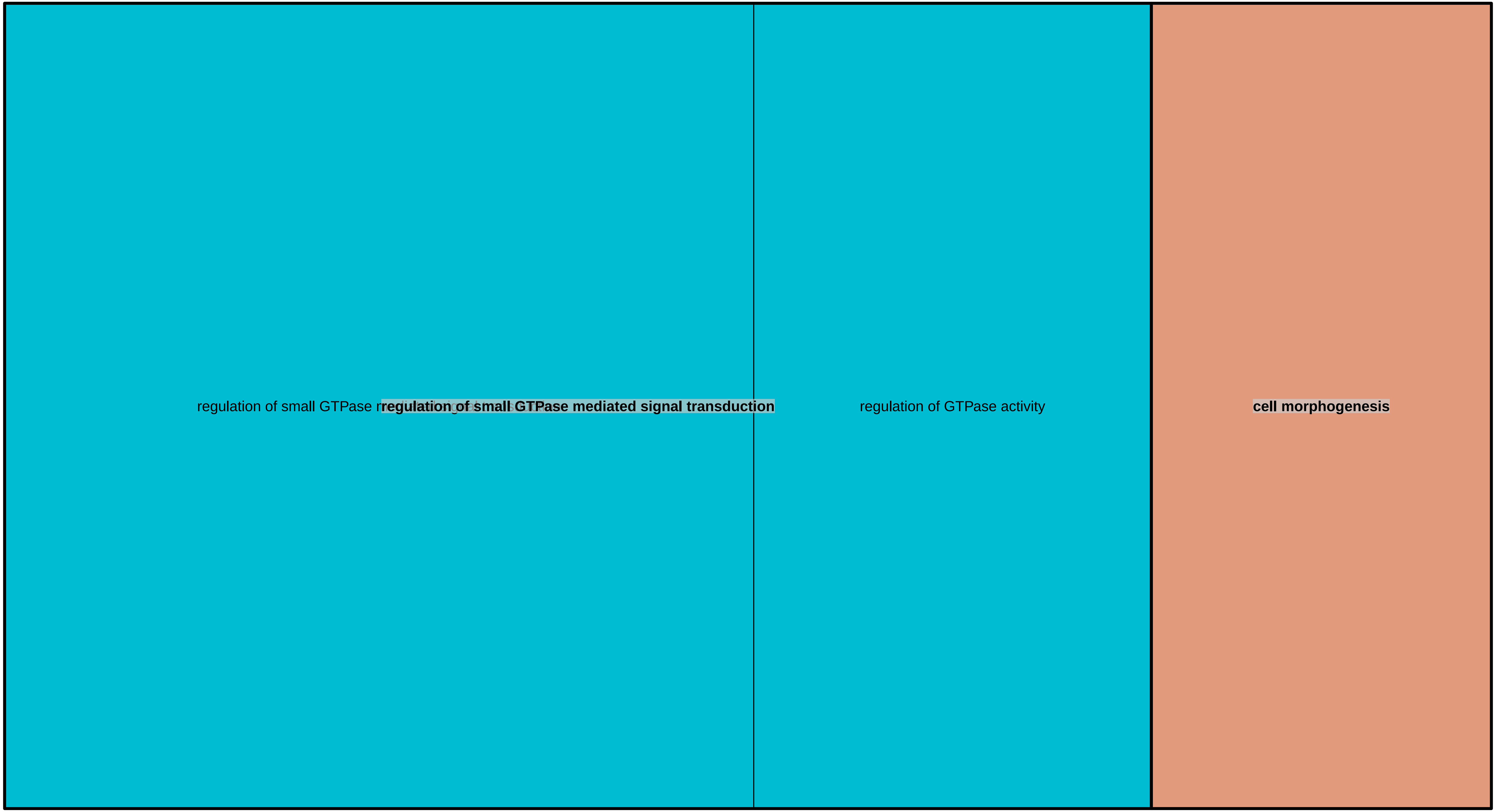

Esophagus − Mucosa Gene Ontology treemap

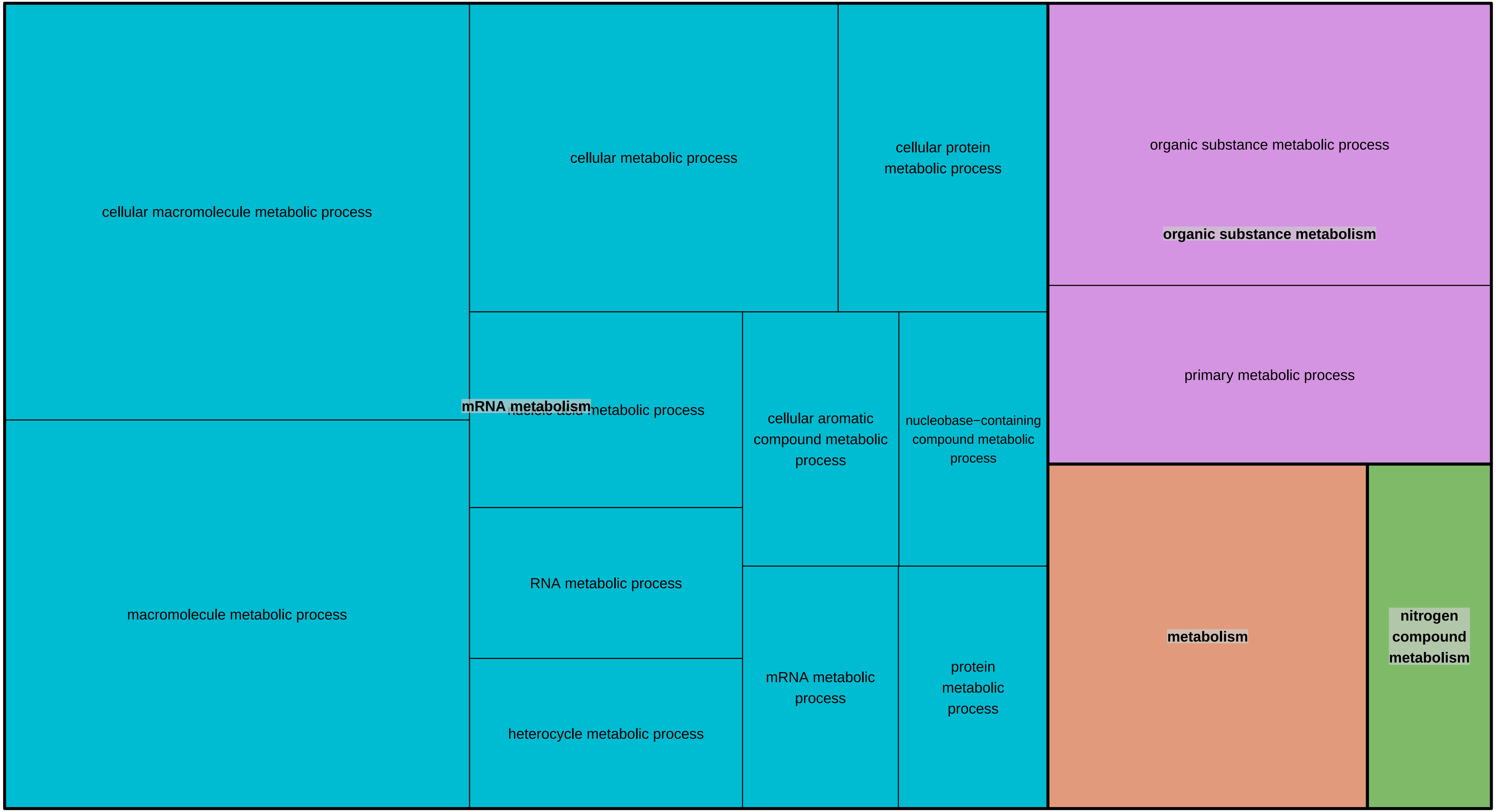

Heart − Left Ventricle Gene Ontology treemap

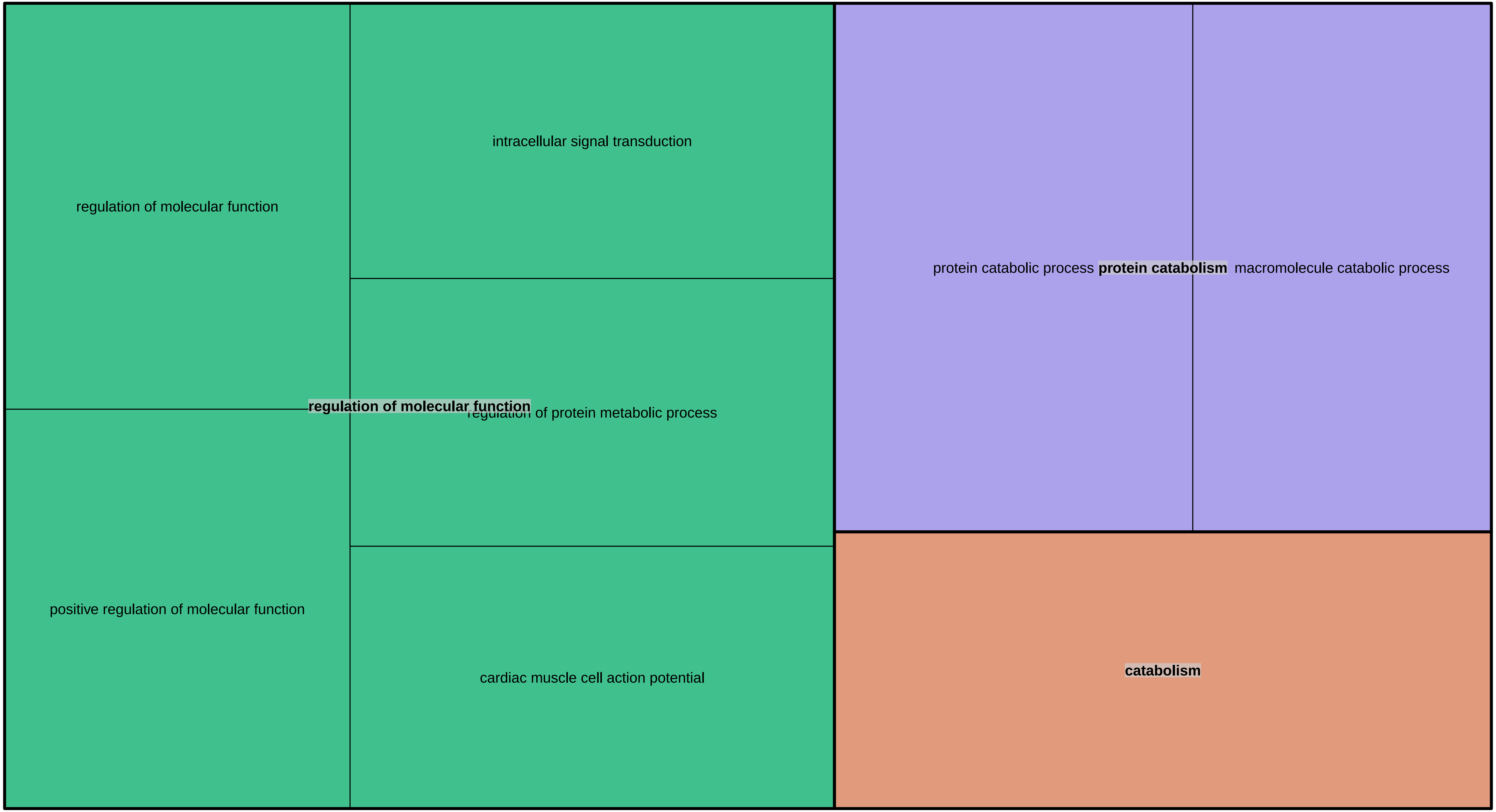

Liver Gene Ontology treemap

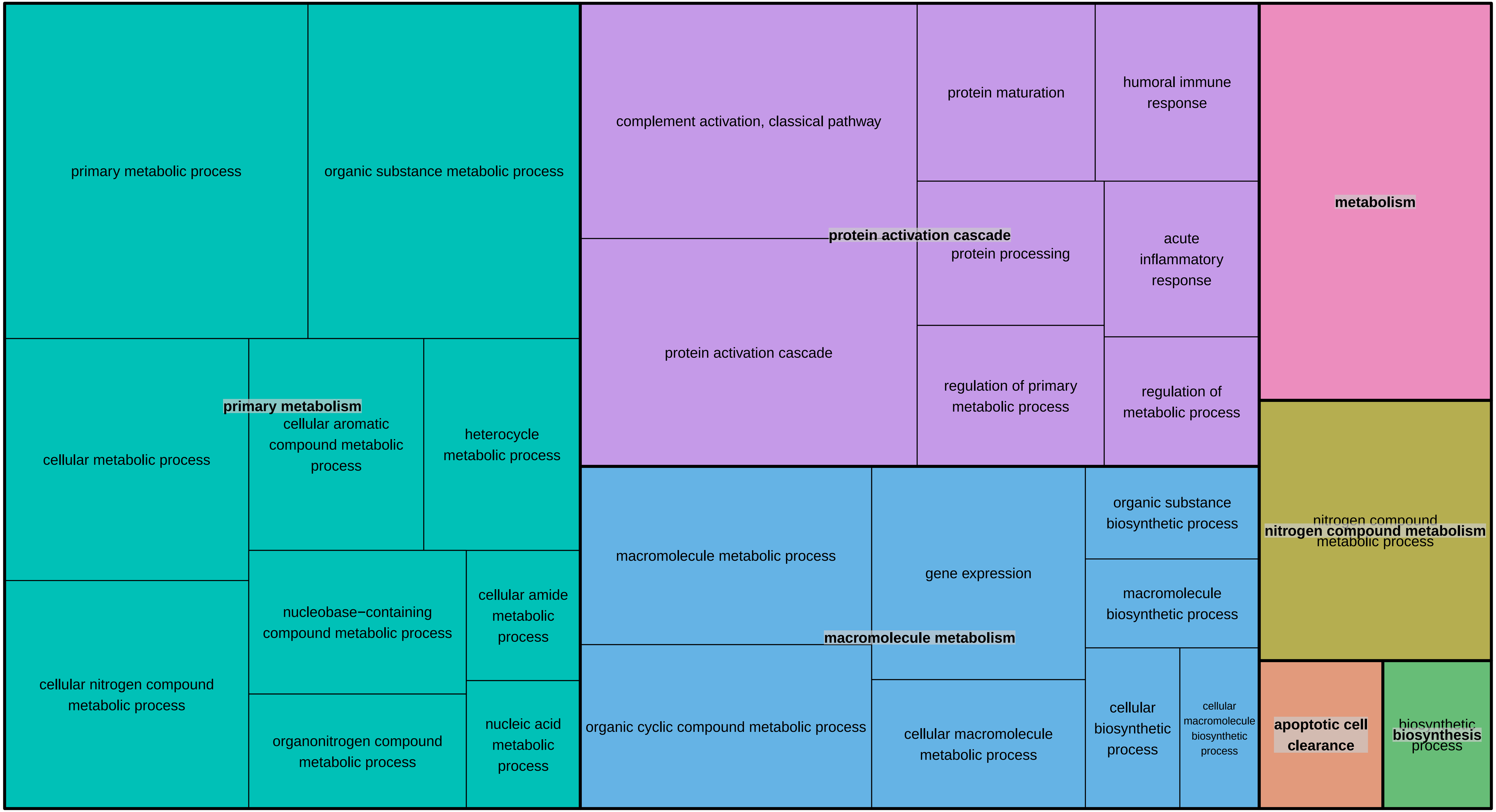

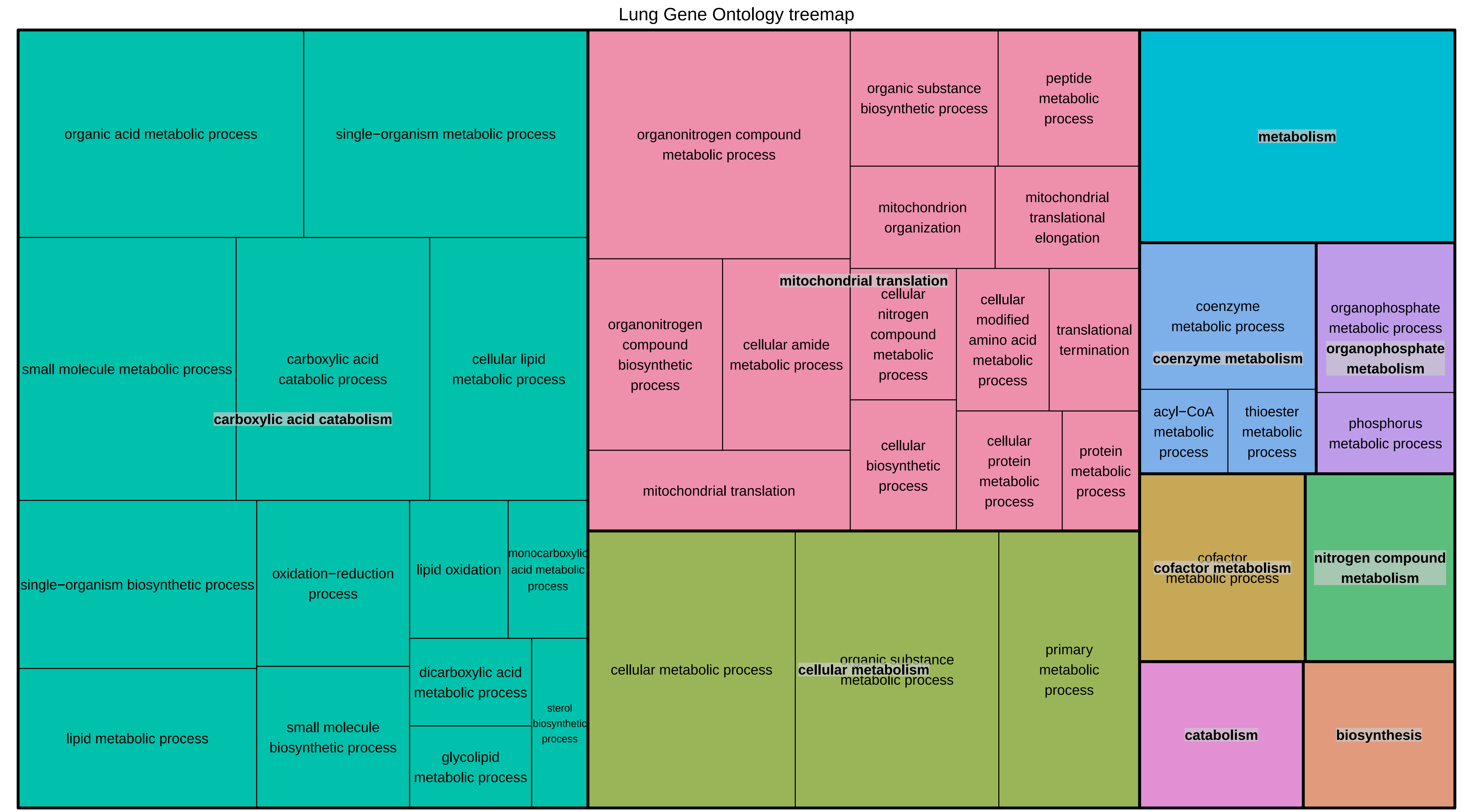

Lung Gene Ontology treemap
organic acid metabolic process
single−organism metabolic process
small molecule metabolic process
carboxylic acid catabolic process
cellular lipid metabolic process
carboxylic acid catabolism
single−organism biosynthetic process
oxidation−reduction process
lipid oxidation
monocarboxylic acid metabolic process
dicarboxylic acid metabolic process
sterol biosynthetic process
glycolipid metabolic process
lipid metabolic process
small molecule biosynthetic process
organonitrogen compound metabolic process
organic substance biosynthetic process
peptide metabolic process
mitochondrion organization
mitochondrial translational elongation
mitochondrial translation
organonitrogen compound biosynthetic process
cellular amide metabolic process
cellular nitrogen compound metabolic process
cellular modified amino acid metabolic process
translational termination
mitochondrial translation
cellular biosynthetic process
cellular protein metabolic process
protein metabolic process
cellular metabolic process
organic substance metabolic process
cellular metabolism
primary metabolic process
metabolism
coenzyme metabolic process
coenzyme metabolism
organophosphate metabolic process
organophosphate metabolism
acyl−CoA metabolic process
thioester metabolic process
phosphorus metabolic process
cofactor metabolic process
cofactor metabolism
nitrogen compound metabolism
catabolism
biosynthesis

#### Muscle − Skeletal Gene Ontology treemap

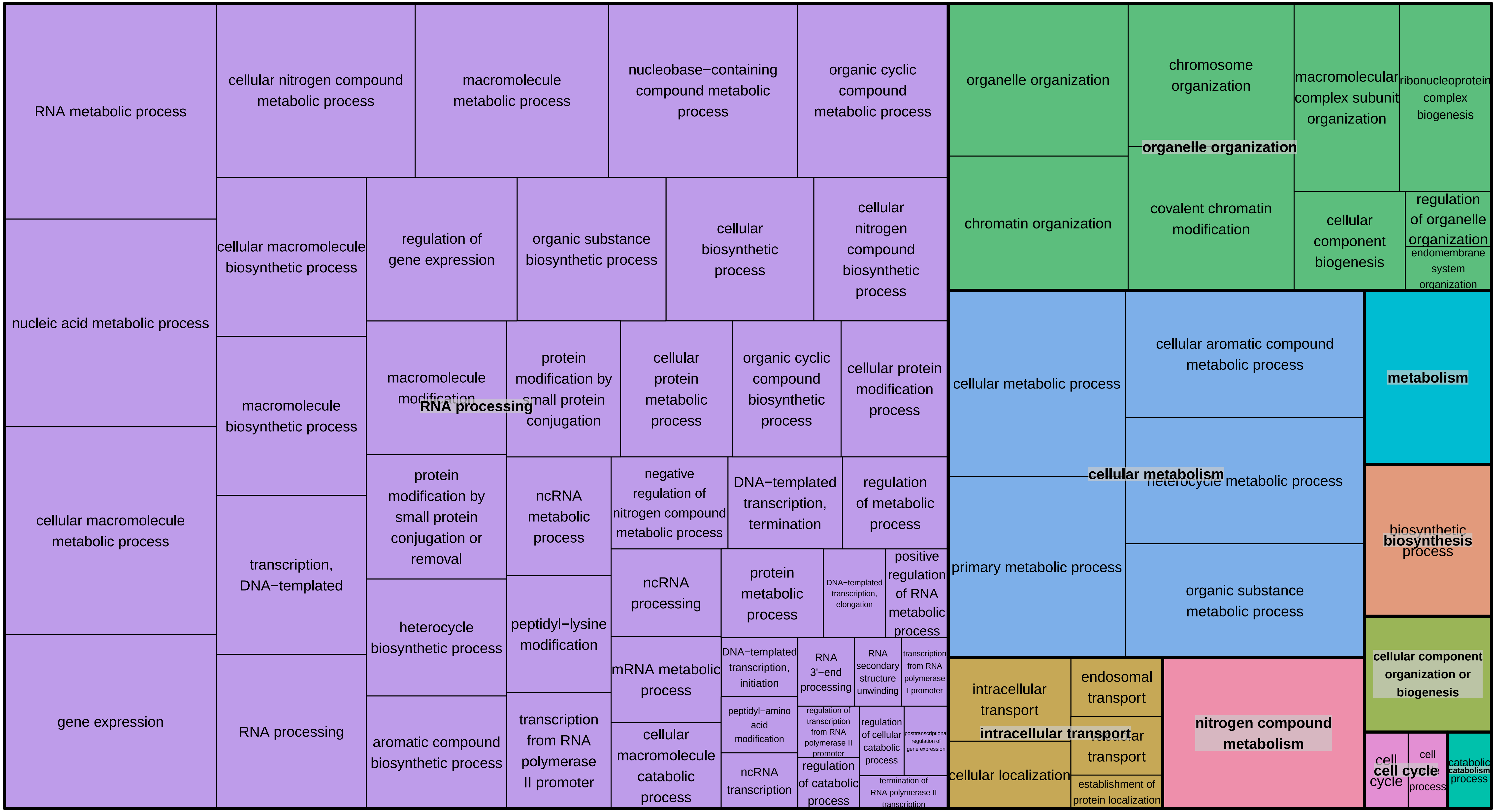

Nerve − Tibial Gene Ontology treemap

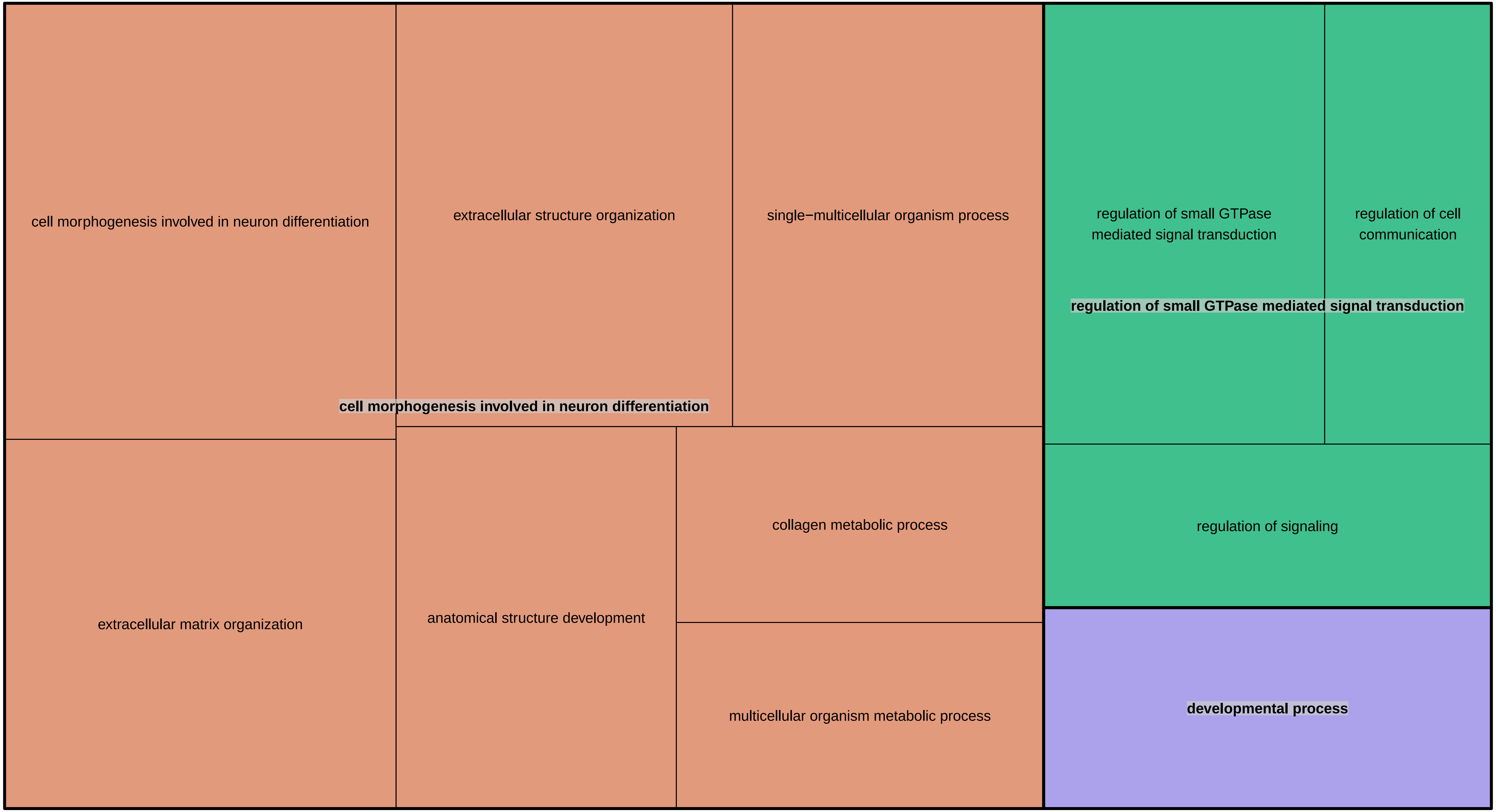

Ovary Gene Ontology treemap

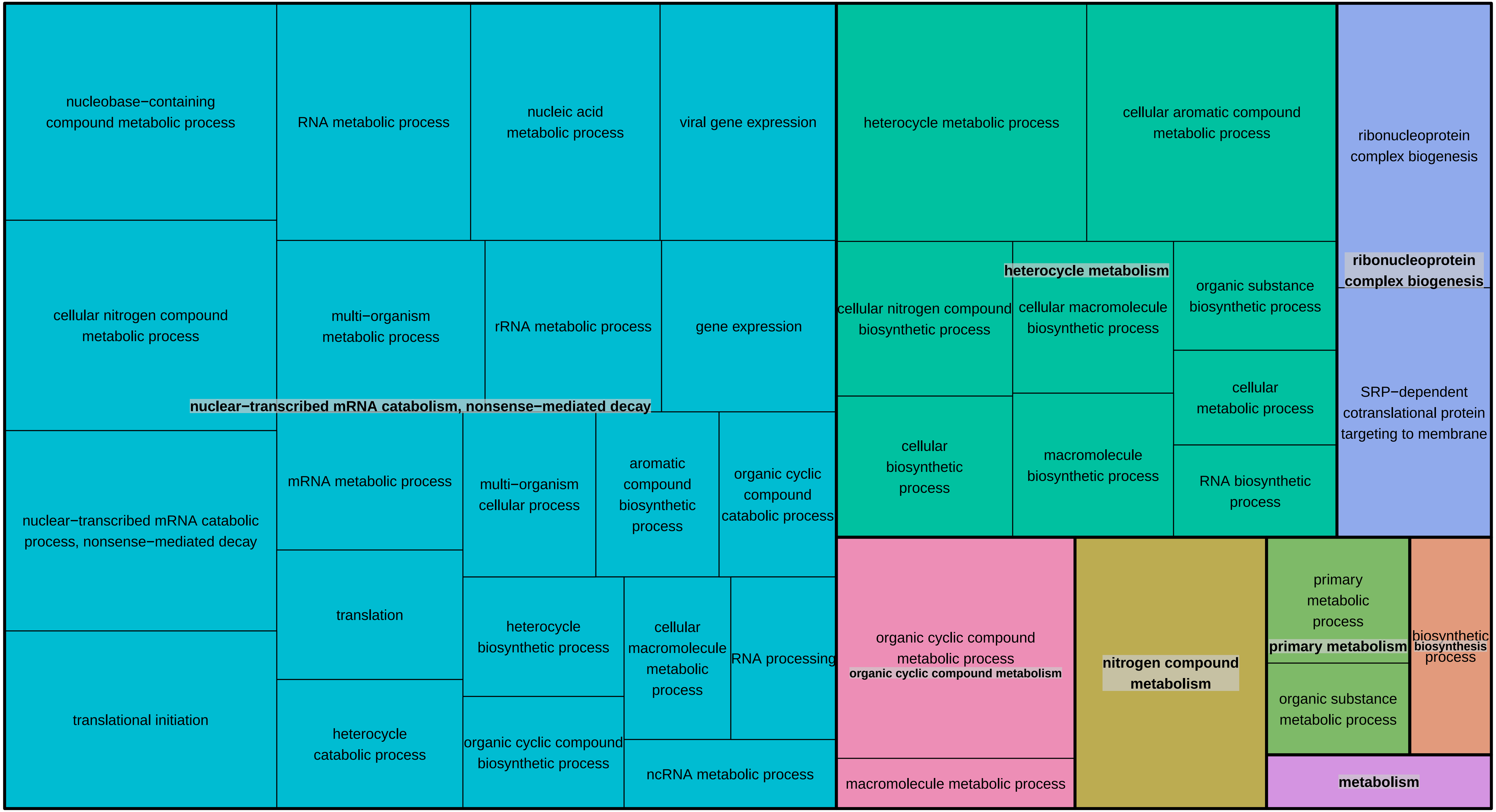

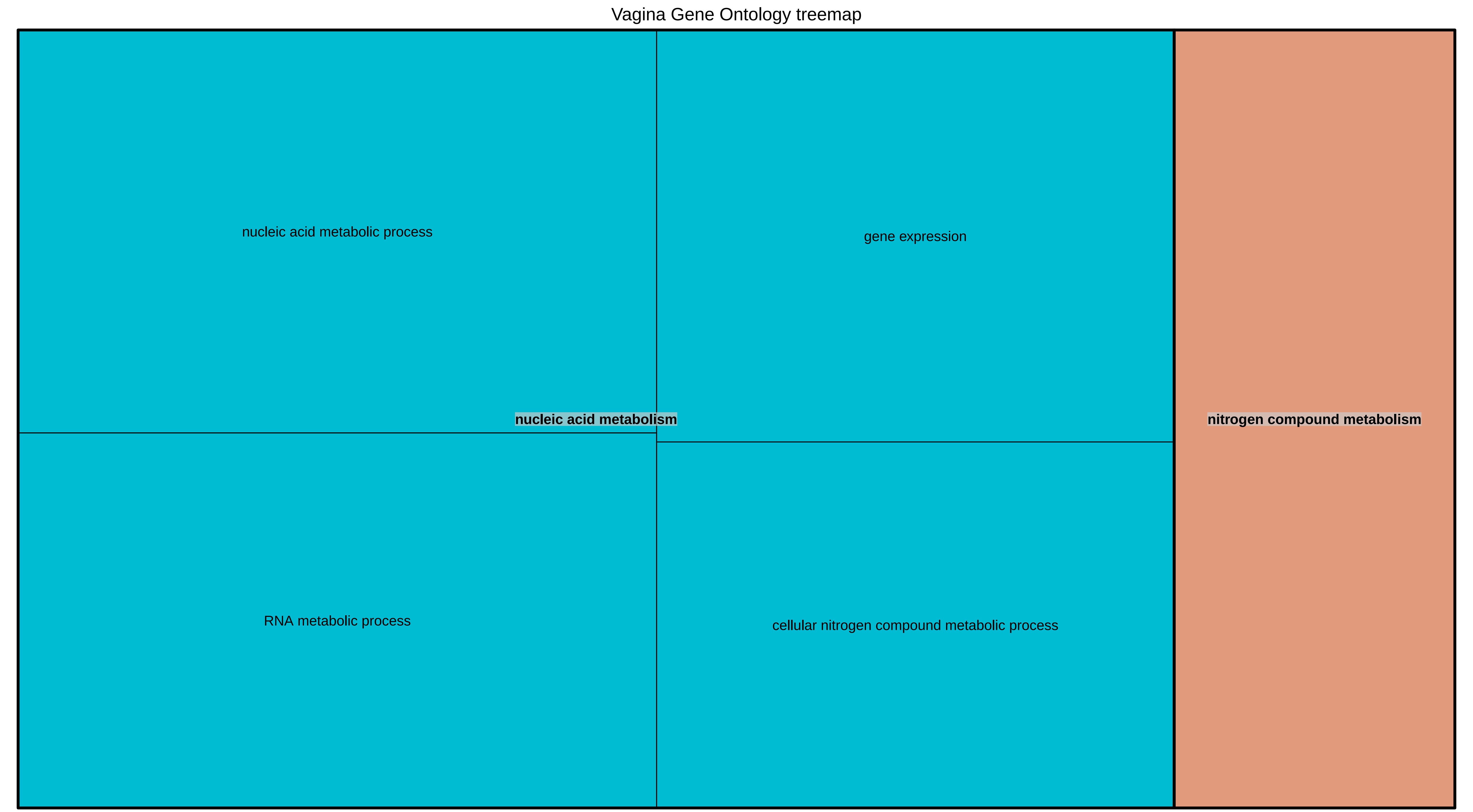
Vagina Gene Ontology treemap
nucleic acid metabolic process
gene expression
nucleic acid metabolism
nitrogen compound metabolism
RNA metabolic process
cellular nitrogen compound metabolic process
