## Supplemental Results for "Predicting tissue-specific gene expression from whole blood transcriptome"

**Supplementary Results and Notes**

#### 1. Prediction accuracy using the base model M1: "WBGE+CF"

For each of the 17,031 genes, in each of the 32 tissues based on the regression model M1 we first assessed whether WBGE significantly informs to TSGE in addition to the demographic covariates (CF), using Log-Likelihood Ratio (LLR) tests (Methods). We found that on average across tissues, for 34% of the genes, WBGE contribute significantly (FDR<=0.05) relative to the CF alone, with up to 67% of the genes in Muscle-Skeletal tissue. For each such gene and tissue we fit the regression base model M1 and estimate the CV accuracies using Pearson Correlation Coefficient (PCC) between the predicted and observed expression across individuals. The mean CV PCC varies from 0.003 in Brain-Caudate (basal ganglia) tissue to 0.289 in Adipose - Visceral (Omentum) tissue, shown in Supplementary Fig. S3a; Supplementary Fig. S3b shows the corresponding plots for all genes. Supplementary Table S2 shows the number of genes having LLR FDR <=0.05 with accuracy above various PCC thresholds. For instance, on average 30.2% of all genes having LLR FDR <= 0.5 have cross-validation PCC >= 0.3. Overall, our results suggest that in many tissues, for a sizeable fraction of genes, an individual's TSGE can be inferred, with some accuracy, from the individual's WBGE.

#### 2. Comparison of models M2 and M1

Overall across all genes and tissues, as expected, M2 yields a greater CV prediction accuracy than M1 (paired one-sided Wilcoxon test p-value = 4.7e-10). More directly, we compared for each gene in each tissue the model accuracy of M2 with that of M1 via LLR test and found that on average across tissues, for 43.2% of the genes, WBSp makes significant additional contribution (FDR <= 0.05), up to 70.7% for Muscle-Skeletal tissue.

#### 3. Contribution of SNPs toward predicting TSGE (M3 model)

Previous eQTL studies have revealed SNPs associated with TSGE in various tissues (Aguet et al., 2017b; GTEx Consortium, 2015). The tool PrediXcan (Gamazon et al., 2015) utilizes the identified eSNPs to predict TSGE. An obvious limitation of the SNP-based TSGE prediction is its scope – it can only be applied to a relatively small subset of genes (10.4% on average per tissue based on (Gamazon et al., 2015)) whose expression variability is significantly associated with genetic variability, i.e., those with detected eSNPs. For such subsets of genes in each tissue, we assessed the extent to which including a gene's eSNPs to the M2 model improves its TSGE prediction. Toward this, we implemented model M3 that includes CF, WBGE, WBSp, and the eSNPs (Methods). Following convention (Guo et al., 2018), we considered SNPs within 1MB of each gene. For each gene we predicted the eSNPs in CV manner, and the top 5 PCs of these eSNPs were then used along with the WBGE and WBSp to predict TSGE in the test set of samples. The prediction accuracies for the genes having at least one eSNPs are summarized in Supplementary Fig. S4, along with the CV accuracy using M2, revealing substantially greater accuracy for M3 relative to M2. Based on LLR test between M3 and M2, we found that, on average for about 60.4% of the genes having at least one eSNP, the eSNPs makes significant additional contribution toward predicting TSGE. The fraction of such genes in each tissue, as well as CV prediction accuracies for those genes for M3 and M2 models are shown in Supplementary Fig. S5.

**4. Comparing model M3 with a SNP-only model**

Next, we directly compared the relative efficacy of SNPs (alone) versus the WBT (model M2) in predicting TSGE. We implemented an additional model M4 involving only the eSNPs (Methods). As for M3 above, eSNPs were estimated from the training set. Our model M4 is qualitatively comparable to PrediXcan (Gamazon et al., 2015) in that they both rely only on eSNPs to predict TSGE. Accuracy of model M4 is generally consistent with those reported for PrediXcan, and the slight differences likely reflect the different versions of the data sets used. For the tissues in common in our study and in (Gamazon et al., 2015), Supplementary Table S3 shows the CV prediction accuracies as reported in (Gamazon et al., 2015) and for our models M4, as well as M2, and M3, only considering the genes having at least one eSNP. Direct comparisons of M4 with M2 and M3 suggest that (i) WBT is a better predictor of TSGE than eSNPs alone, but (ii) eSNPs together with WBT performs better than WBT alone.

**5. Genes having eSNPs have greater predictability**

For a SNP-based model such as M4 and (J. Wang et al., 2016), TSGE prediction accuracy is expected to be greater for genes with greater number of eSNPs. Interestingly, however, a previous work predicting lung expression based on Whole blood expression found the prediction accuracy to be greater for genes with greater number of eSNPs, despite not using eSNPs in the model (Halloran et al., 2015). To assess generality of this previous finding, we defined three subsets of genes: those with no reported eSNPs, those having 1-4 eSNPs, and those having >= 5 eSNPs. Supplementary Fig. S5 shows that our M2 model (which notably does not include eSNPs) can predict TSGE with a greater accuracy for the genes having eSNPs relative to those that do not. This suggests that eSNPs may inform their target gene's expression indirectly through mechanisms involving other genes' expression, which is captured by M2. This result is also consistent with the fact the genes with greater connectivity (potential regulators) have greater predictability.

### Supplementary Figures:

**Supplementary Fig S1:** Bar plot showing the number of common samples (individuals) present between a tissue and Whole Blood for all the tissues available in GTEx.

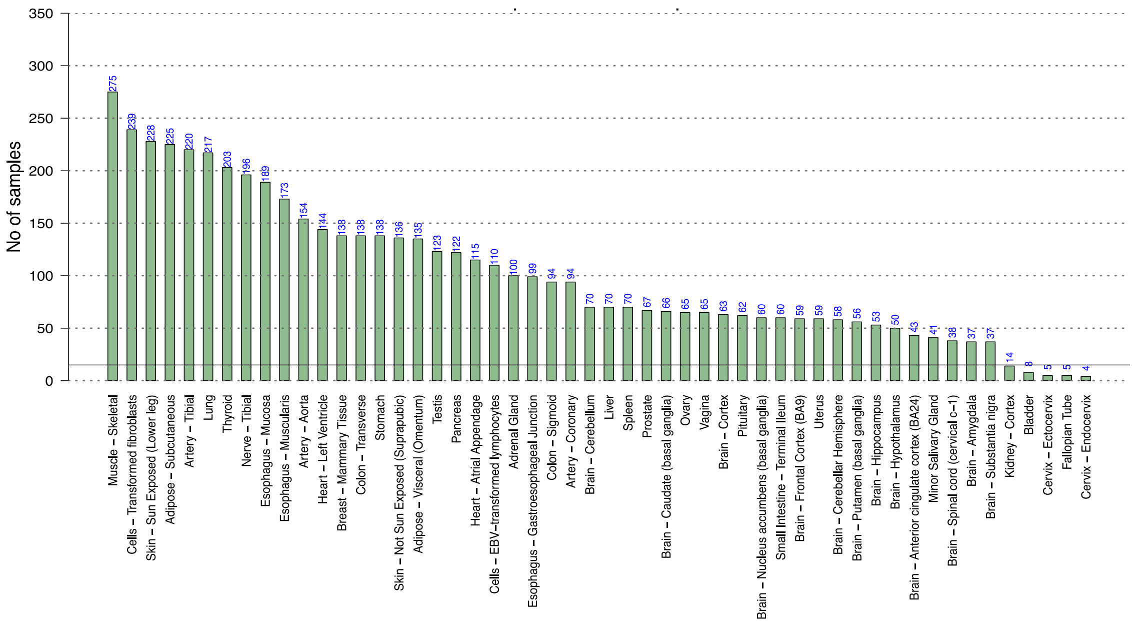


**Supplementary Fig S2:** Prediction accuracy of gene expression in target tissue using model M2 (WBGE+WBSp+CF). The prediction accuracy in terms of PCC for all the genes are plotted here. The blue points indicate the mean.

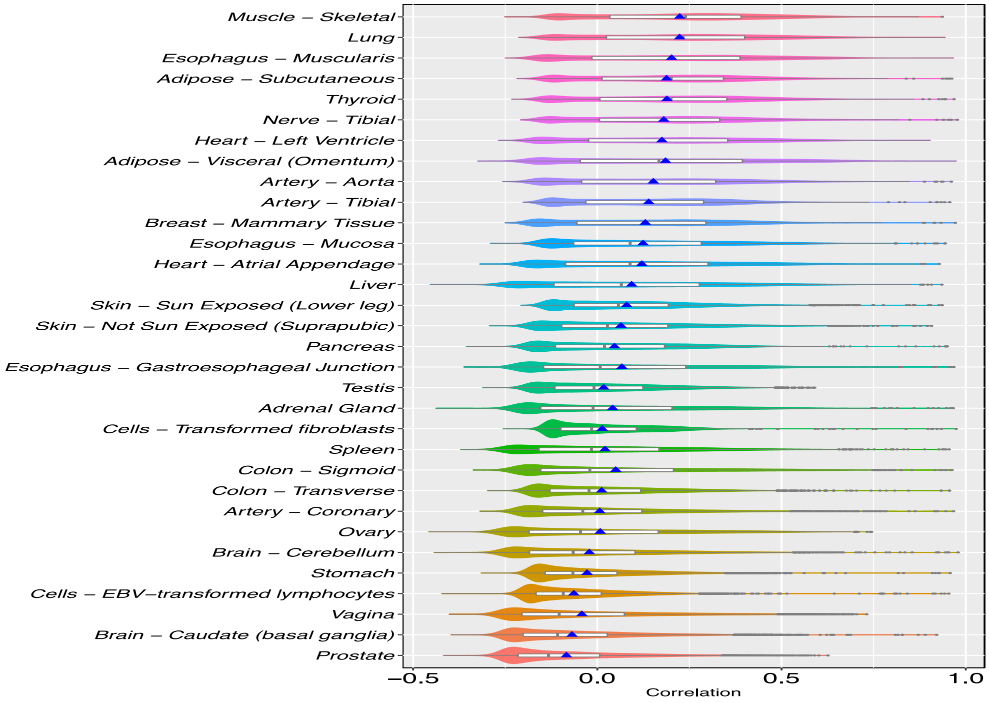

**Supplementary Fig S3 (a):** Prediction accuracy of gene expression in target tissue using model M1 (WBGE+CF) in cross-validation manner (with 5 folds and 25 independent runs) is calculated and expressed in terms of Pearson correlation coefficient (PCC). The prediction accuracy for only those genes with LLR (between M1 and CF) FDR <= 0.05 are plotted here. The blue points indicate the mean accuracy, and the numbers beside each of the violin plots denotes the percentage of genes which has LLR FDR<=0.05 among all the genes in this study. **(b)** The prediction accuracy in terms of PCC for all the genes are plotted here. The blue points indicate the mean.

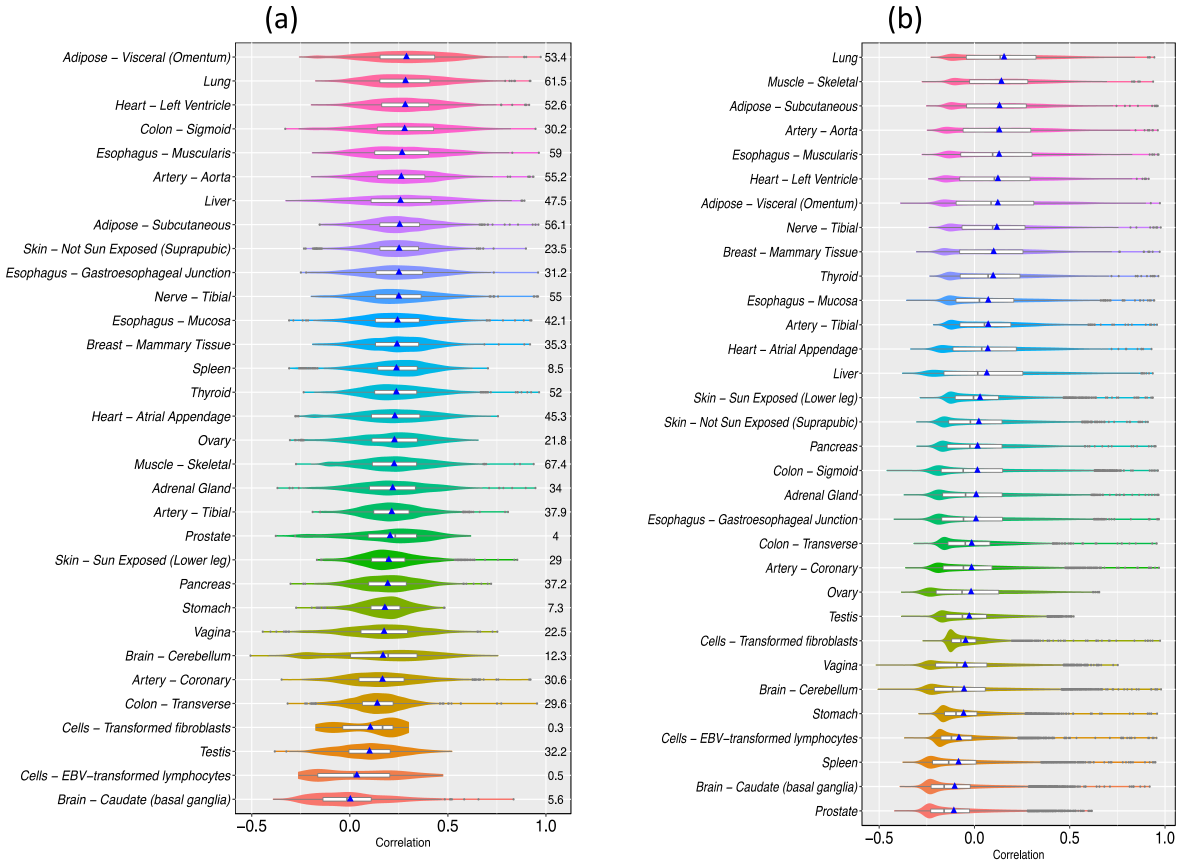

**Supplementary Fig S4**: The prediction scores obtained from model M1 (WBGE+CF), M2 (WBGE+WBSp+CF) and M3 (WBGE+WBSp+eSNPs+CF) for the genes with at least 1 eSNP are shown in the boxplots. The number of genes with at least 1 eSNPs for each tissue is shown in the top.

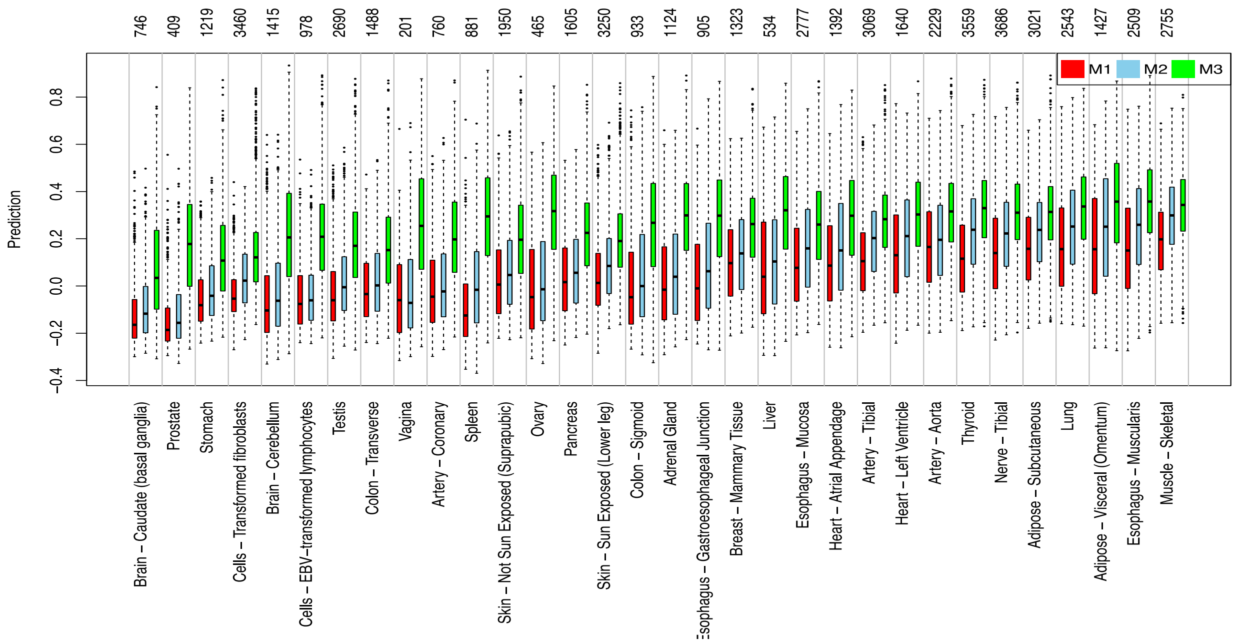


**Supplementary Fig S5**: For the genes having significant contribution from eSNPs over M2 for its expression prediction, comparison of its prediction accuracies obtained from model M1 (WBGE+CF), M2 (WBGE+WBSp+CF) and M3 (WBGE+WBSp+eSNPs+CF) are shown. In the top row the number of genes having significant contribution from eSNPs are shown, and in the parenthesis the number of genes having at least one eSNPs are shown.

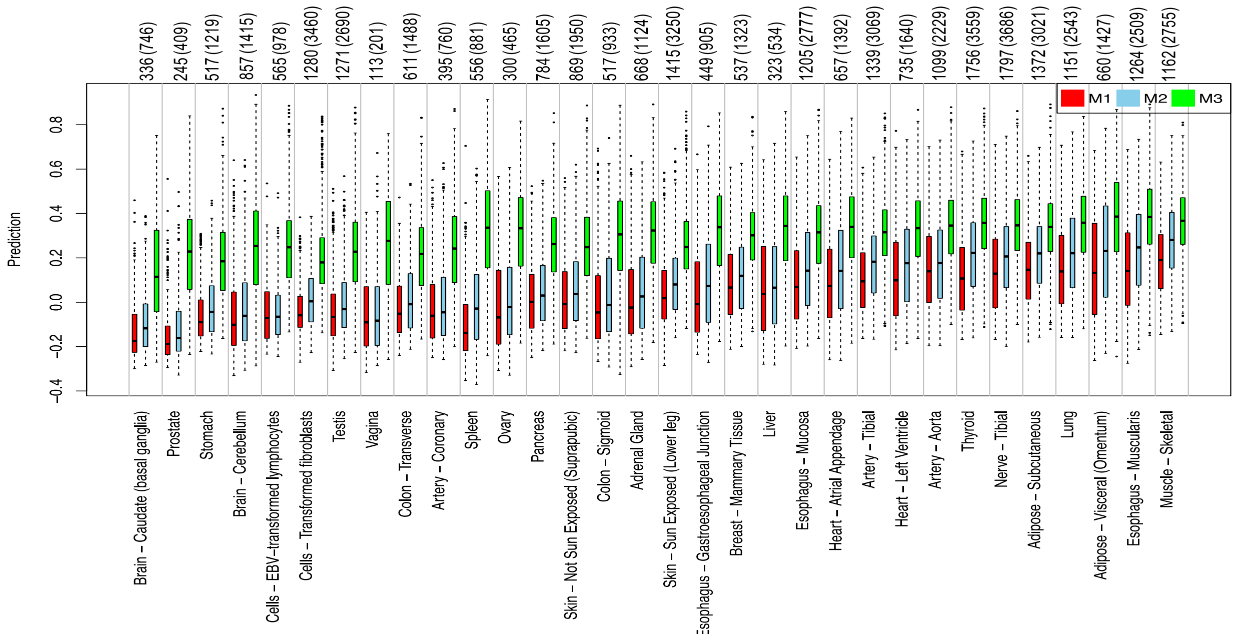

**Supplementary Fig S6**: Prediction accuracies of the genes calculated using model M2 for genes with 0, 1-4, and >4 eSNPs, showing increased prediction accuracies for expression of the genes having at least one eSNPs than those without eSNPs.

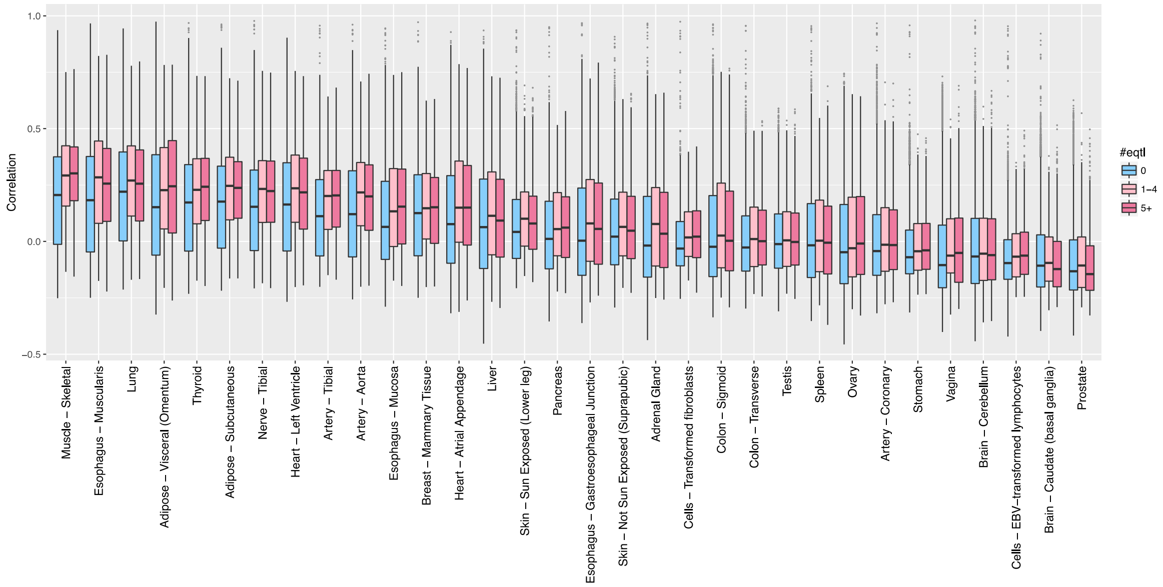


**Supplementary Fig S7**: Plot shows the overlap among the top 25 percentile (based on CV PCC) of tissue-specific predictable (FDR LLR(M2~CF) <=0.05) genes in all tissue's pairs in terms of Jaccard index.

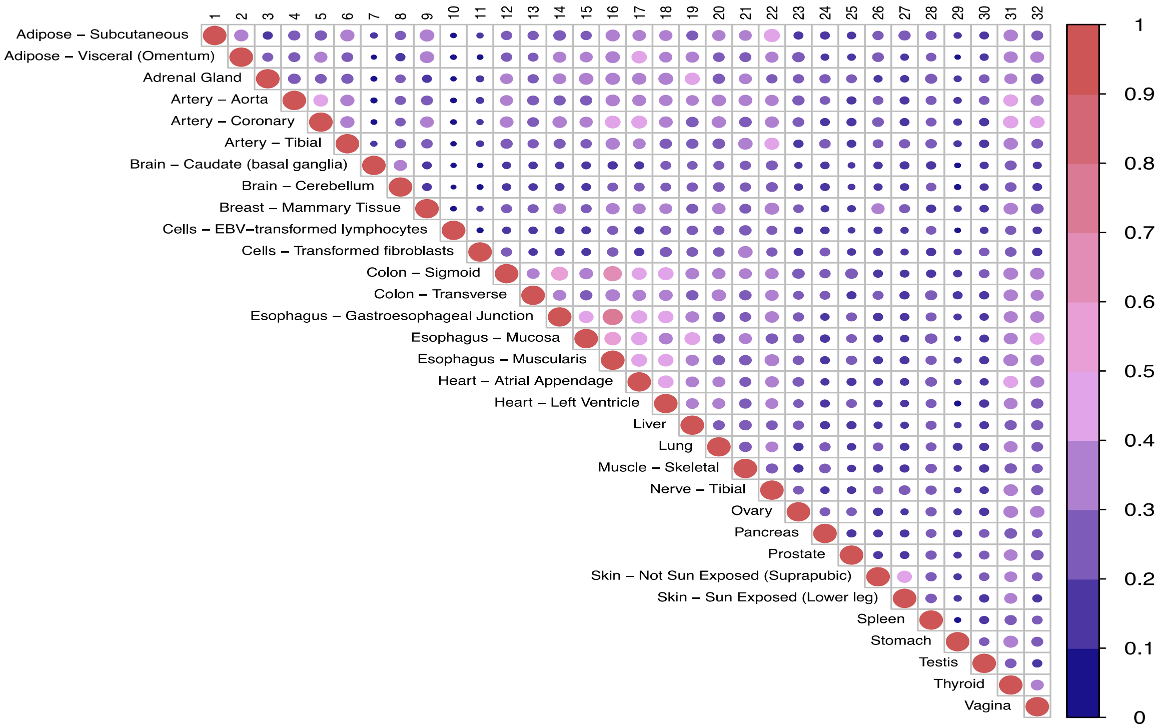

**Supplementary Fig S8. (a)** The figure shows the broad biological processes that are significantly enriched among the highly predictable genes in each Tissue. Only the tissues with at least 5 enriched terms are included. **(b)** Combined view of processes enriched in any of the tissues. [**Separate Files: Basu-Imputation-FigS8a.pdf, Basu-Imputation-FigS8b.pdf]**

**Supplementary Fig S9**: Bar plot showing the fraction of housekeeping and tissues-specific among TSPG (see text), and overall.

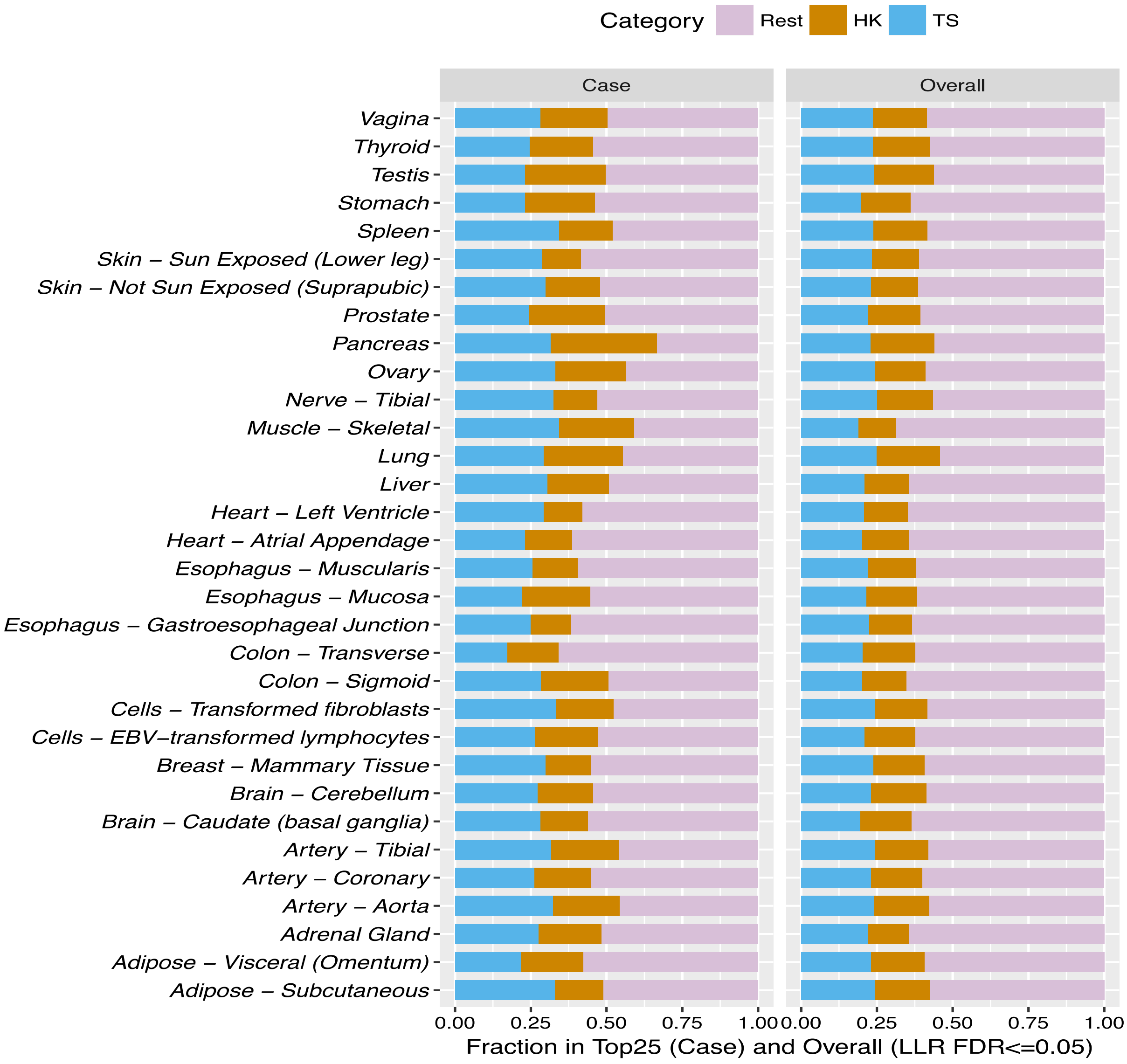

**Supplementary Figure S10.** For 23 disease-tissues pairs the figure shows disease prediction accuracy score (in terms of auROC) using observed TSGE, predicted TSGE and WBGE of all the genes having LLR FDR <= 0.05 and PCC >= 0.3, in 5-fold CV scenario across 50 independent runs. See text for the description of 23 disease-tissues shown.

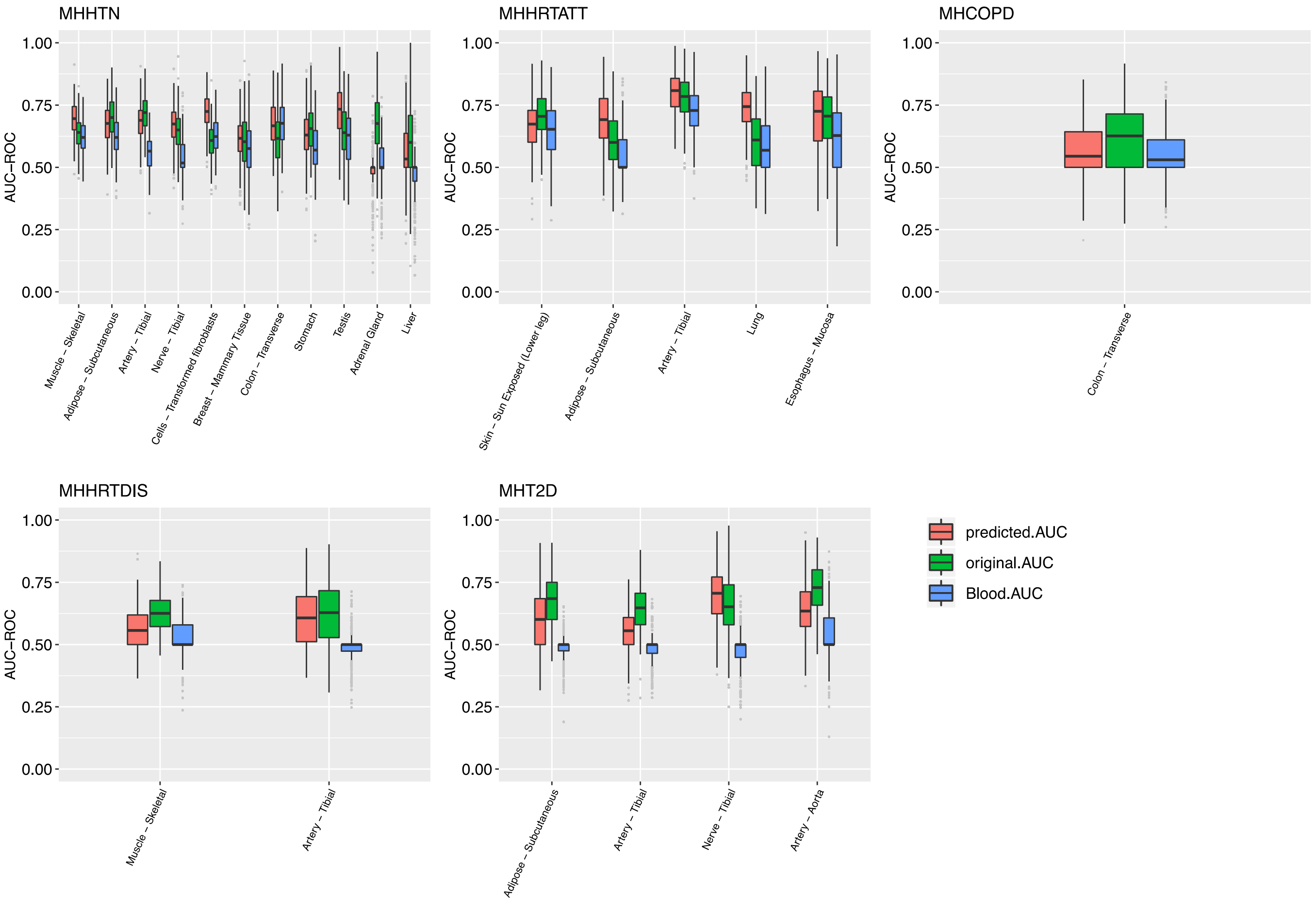
